# Epigenetic changes regulating the epithelial-mesenchymal transition in human trophoblast differentiation

**DOI:** 10.1101/2024.07.02.601748

**Authors:** William E. Ackerman, Mauricio M. Rigo, Sonia C. DaSilva-Arnold, Catherine Do, Mariam Tariq, Martha Salas, Angelica Castano, Stacy Zamudio, Benjamin Tycko, Nicholas P. Illsley

**Affiliations:** Department of Obstetrics and Gynecology and AI.Health4All Center for Health Equity Using Machine Learning and Artificial Intelligence, University of Illinois College of Medicine, Chicago, USA; Hackensack Meridian Health Center for Discovery and Innovation, Nutley, NJ; Department of Obstetrics and Gynecology, Hackensack University Medical Center, Hackensack NJ

**Keywords:** Trophoblast, differentiation, epithelial-mesenchymal transition, methylation

## Abstract

The phenotype of human placental extravillous trophoblast (EVT) at the end of pregnancy reflects both first trimester differentiation from villous cytotrophoblast (CTB) and later gestational changes, including loss of proliferative and invasive capacity. Invasion abnormalities are central to two major placental pathologies, preeclampsia and placenta accreta spectrum, so characterization of the corresponding normal processes is crucial. In this report, our gene expression analysis, using purified human CTB and EVT cells, highlights an epithelial-mesenchymal transition (EMT) mechanism underlying CTB-EVT differentiation and provides a trophoblast-specific EMT signature. In parallel, DNA methylation profiling shows that CTB cells, already hypomethylated relative to non-trophoblast cell lineages, show further genome-wide hypomethylation in the transition to EVT. However, a small subgroup of genes undergoes gains of methylation (GOM) in their regulatory regions or gene bodies, associated with differential mRNA expression (DE). Prominent in this GOM-DE group are genes involved in the EMT, including multiple canonical EMT markers and the EMT-linked transcription factor *RUNX1*, for which we demonstrate a functional role in modulating the migratory and invasive capacities of JEG3 trophoblast cells. This analysis of DE associated with locus-specific GOM, together with functional studies of an important GOM-DE gene, highlights epigenetically regulated genes and pathways acting in human EVT differentiation and invasion, with implications for obstetric disorders in which these processes are dysregulated.

## Introduction

Differentiation of cytotrophoblast (CTB) into invasive extravillous trophoblast (EVT) is a crucial element in human placentation. The EVT at the distal ends of chorionic villi attach the placenta to the decidua, acting as anchor points for the placenta (Knofler et al., 2019). EVT cells moving into the uterus from the implantation site early in pregnancy form plugs in the spiral arteries, preserving the physiologically hypoxic environment around the conceptus and promoting trophoblast proliferation in the rapidly growing placenta (Weiss et al., 2016). EVT are a key cell type in remodeling the maternal spiral arteries to efficiently deliver maternal blood to the intervillous space (Knofler et al., 2019).

Elucidation of the differentiation and invasion processes, whereby anchored CTB are transformed into invasive EVT, is central to our understanding of placental development and major obstetric conditions involving this organ. One such condition is preeclampsia (PE), in which EVT invasion is more limited than in normal pregnancy, leading to decreased spiral artery remodeling, reduced uteroplacental blood flow, placental hypoxia, maternal endothelial damage and the clinical sequelae (Phipps et al., 2019). At the other extreme is placenta accreta spectrum (PAS), where EVT invade more deeply into the uterus, supplanting decidual and myometrial tissue. This results in atypical remodeling of deeper uterine vessels, possibly as deep as the radial arteries (Tantbirojn et al., 2008), leading to potentially fatal hemorrhage when PAS goes undetected. In both disorders, the invasion by EVT is abnormal, raising the question of the control mechanisms.

Prior research has suggested that a key molecular mechanism governing trophoblast differentiation and invasion is the epithelial-mesenchymal transition (EMT; (Davies et al., 2016; Kokkinos et al., 2010; Vicovac and Aplin, 1996)). The EMT is common to multiple processes such as gastrulation, wound healing and cancer metastasis whereby anchored epithelial cells are converted to a more invasive, mesenchymal phenotype (Kalluri and Weinberg, 2009). We have demonstrated changes in gene expression indicative of an EMT in both first and third trimester EVT (DaSilva-Arnold et al., 2015; DaSilva-Arnold et al., 2018). While first trimester EVT occupy a relatively mesenchymal position on the EMT spectrum, term EVT show a degree of regression towards the epithelial pole, consistent with the loss of invasiveness observed in mid- and late pregnancy (Illsley et al., 2020).These results prompted us to investigate the differentiation process in more depth, via RNA-sequencing, to confirm the role of the EMT and to develop a term trophoblast EMT signature. Further, we investigate the role of DNA methylation which, based on prior reports, likely plays a role in regulating human trophoblast differentiation (Anton et al., 2014; Chen et al., 2013; Nordor et al., 2017; Zhang et al., 2021).

## Results

### Comparison of CTB and EVT transcriptomes highlights differential expression of genes involved in trophoblast differentiation and the EMT

Placental tissue samples were obtained from nine normal, term pregnancies. Clinical data are in **Supplementary Table S1**. All nine placentas had CTB and EVT purified from placental basal plate. In four of the nine placentas CTB from the chorionic villous core of the placenta (vCTB) were also obtained and analyzed. Illumina EPIC (800K) DNA methylation assays were performed on the same subset of four samples. There were no differences in the clinical parameters between the full set of samples and the EPIC/vCTB subset. For each placental sample, CTB and EVT were obtained from the same tissue and are thus “paired”. The vCTB extracted from chorionic villous tissue samples and the DNA utilized for the methylation assays were obtained from the same samples from which CTB and EVT were isolated for RNA preparation.

Comparison between the RNA-sequencing of the nine sets of paired CTB and EVT yielded > 23,000 transcripts. Paired analysis, using the threshold parameters of (1) FDR threshold of ≤ 0.05, (2) featureCount (baseMean) ≥ 10 and (3) a fold change in expression of ≥ 1.5, restricted differential expression (DE) to ∼7600 transcripts. This comprised 3817 up-regulated genes and 3772 down-regulated genes as shown in the volcano plot in **Figure 1A**. Multidimensional visualization via principal components analysis of DE) between vCTB, CTB, and EVT (**Figure 1B**) shows that the CTB and EVT clusters are well separated. This confirms the expected clear distinction between these cell types, based both on our flow cytometry results (see Materials and Methods) and on gene expression profiling. The marked separation in the first component between CTB and EVT explains >80% of the variance between them. The top 50 leading edge genes (25 with a positive coefficient, 25 with a negative coefficient) driving this separation are shown in **Figure 1C**. Several genes that are expressed primarily in trophoblast, including *HLA-G*, *PAPPA*, *CSH1* (positive) and *PLAC4*, S*IGLEC6*, and *ERVFRD-1* (negative) are prominent. The clear difference between CTB and EVT is further confirmed by the clustergram for the top 1000 genes (**Figure 1D)**. In addition, the vCTB cluster aligns closely with CTB in the PCA plot (**Figure 1B**), suggesting that the phenotypic differences between cytotrophoblast obtained from the basal plate (CTB) and villous core (vCTB) are small relative to those between CTB and EVT. In the clustergram (**Figure 1D**) the vCTB, while differentiated from the EVT, are intermixed with the CTB, indicating that the CTB obtained from the basal plate versus the villous core differ much less than either cell type from EVT.

**Figure 1:**
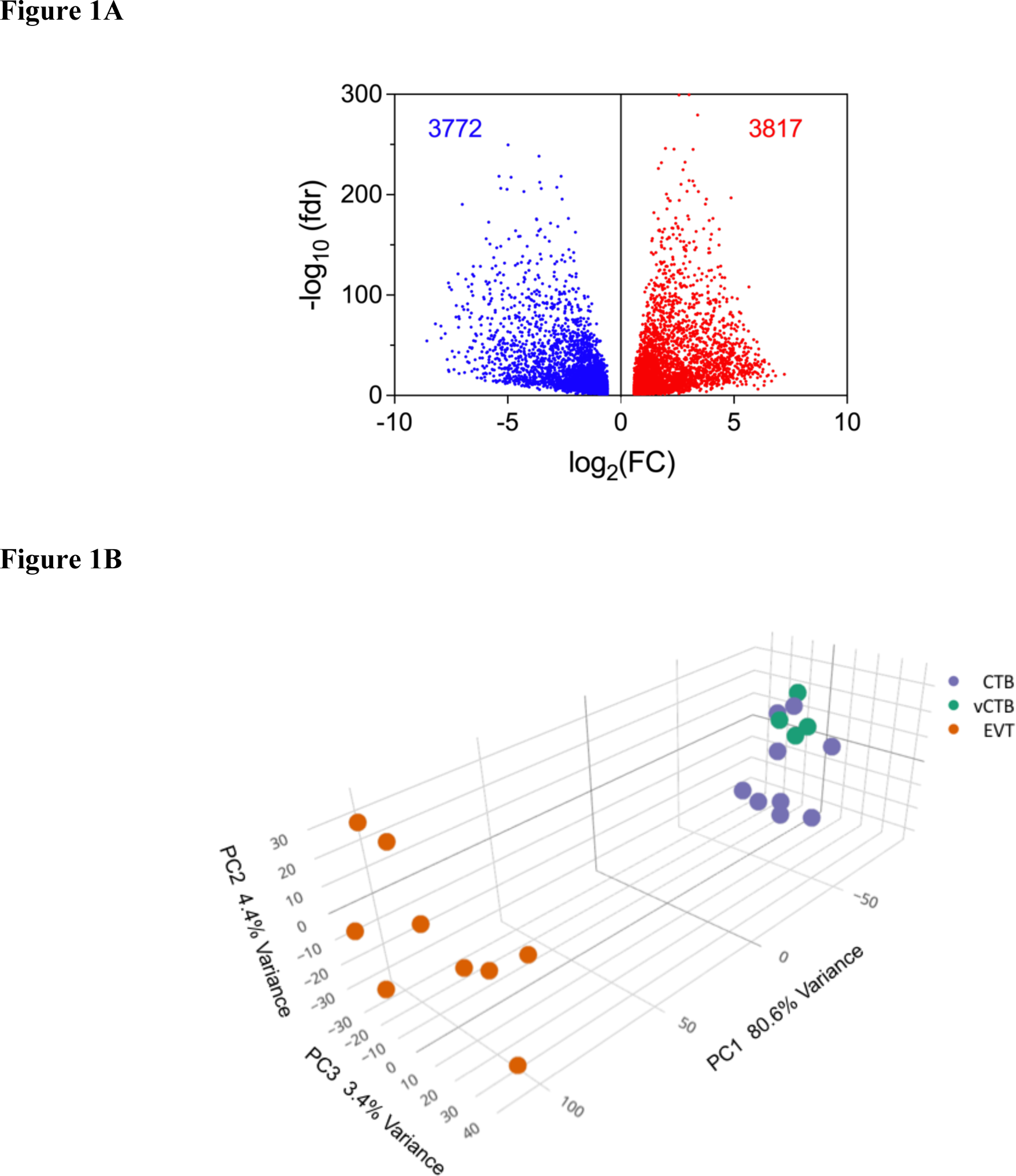
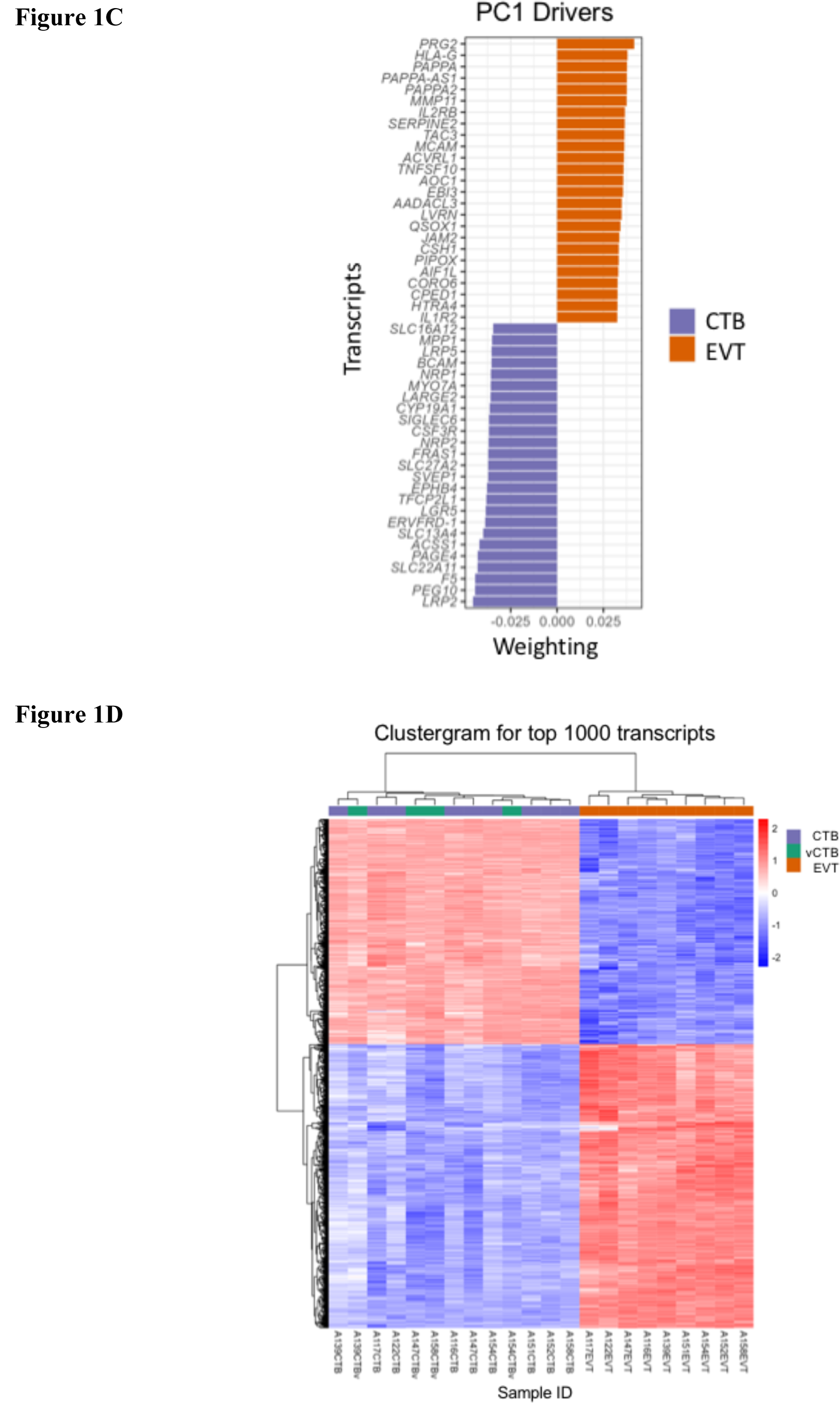
Characteristics of the differentially expressed genes in EVT compared to CTB**. *(A)*** *Volcano plot of RNA-sequencing DE data showing decreased expression (blue) and increased expression (red). Paired analysis, using the threshold parameters of (1) FDR ≤ 0.05, (2) featureCount (baseMean) ≥ 10, (3) a fold change of ≥ 1.5. **(B)** PCA plot of CTB (purple), vCTB (green) and EVT (orange) using the fullDE geneset. **(C)** Top 50 driver genes (25 positive, 25 negative) for the first component of the PCA. **(D)** Clustergram for CTB (purple), vCTB (green) and EVT (orange) for the top 1000 genes*.

### EMT as a part of trophoblast differentiation

The enrichment for EMT as a biological process is consistent with our results obtained previously from the PCR arrays of EMT-associated genes (DaSilva-Arnold et al., 2015; DaSilva-Arnold et al., 2018; Natenzon et al., 2022). We therefore examined the correspondence between the RNA-seq results for the CTB/EVT differentiation process and a library of EMT genes assembled from multiple sources (Cheng et al., 2012; Gamage et al., 2018; Groger et al., 2012; Taube et al., 2010; Zhao et al., 2015). This list comprises 563 genes, derived mostly from cancer studies. In the intersection with our DE data, 345 (>60%) genes overlapped with this EMT gene library (**Supplementary Table S2**). Among those (DE-EMT) genes are many regarded as canonical markers of the EMT, including cell junctional elements (*CLDN1, MARVELD3, OCLN*) cytoskeletal components (*ANK3, COL5A1, KRT19*), extracellular matrix genes (*FN1, ITGA5, ITGB4*), secreted proteases (*ADAM12, ADAM 19, MMP21*) and signaling genes (*BMP7, EGFR, TGFB2*). Notable also are two EMT master regulators, *ZEB1* and *SNAI1,* both of which are up- regulated in EVT. The high proportion of canonical pro-EMT genes showing up-regulated expression in EVT compared to CTB points to the EMT mechanism as a major contributor to EVT differentiation.

### Functional enrichment of the EMT mechanisms in differentiation

Following the identification of genes differentially expressed between CTB and EVT, we determined the functional processes that are enriched among these DE genes. Data generated by DESeq2 was used to test for enrichment of the Hallmark genesets (v7.4) obtained from the Molecular Signatures Database (MSigDB) collections (EVT vs CTB; ≥ 1.5 fold-change ; FDR < 0.05). The extent (or loss) of enrichment for the top 15 processes is quantified in **Table 1A**. This analysis revealed significant enrichment in genes employed as indicators of the EMT (**Table 1B**; baseMean >10, fold change > 1.5, p-adj <0.05). The volcano-bubble plot in **Figure 2A** summarizes the functional enrichment results. It shows most conspicuously a significant enrichment in the EMT geneset, corroborating the role of this mechanism in CTB to EVT differentiation. These results are confirmed in the radar plot of enriched genesets shown in **Figure 2B**. In parallel with up-regulation of the EMT geneset is a reduction in the G2M and E2F target genesets, demonstrating a loss of proliferative capacity in the transformation from CTB to EVT.

**Figure 2:**
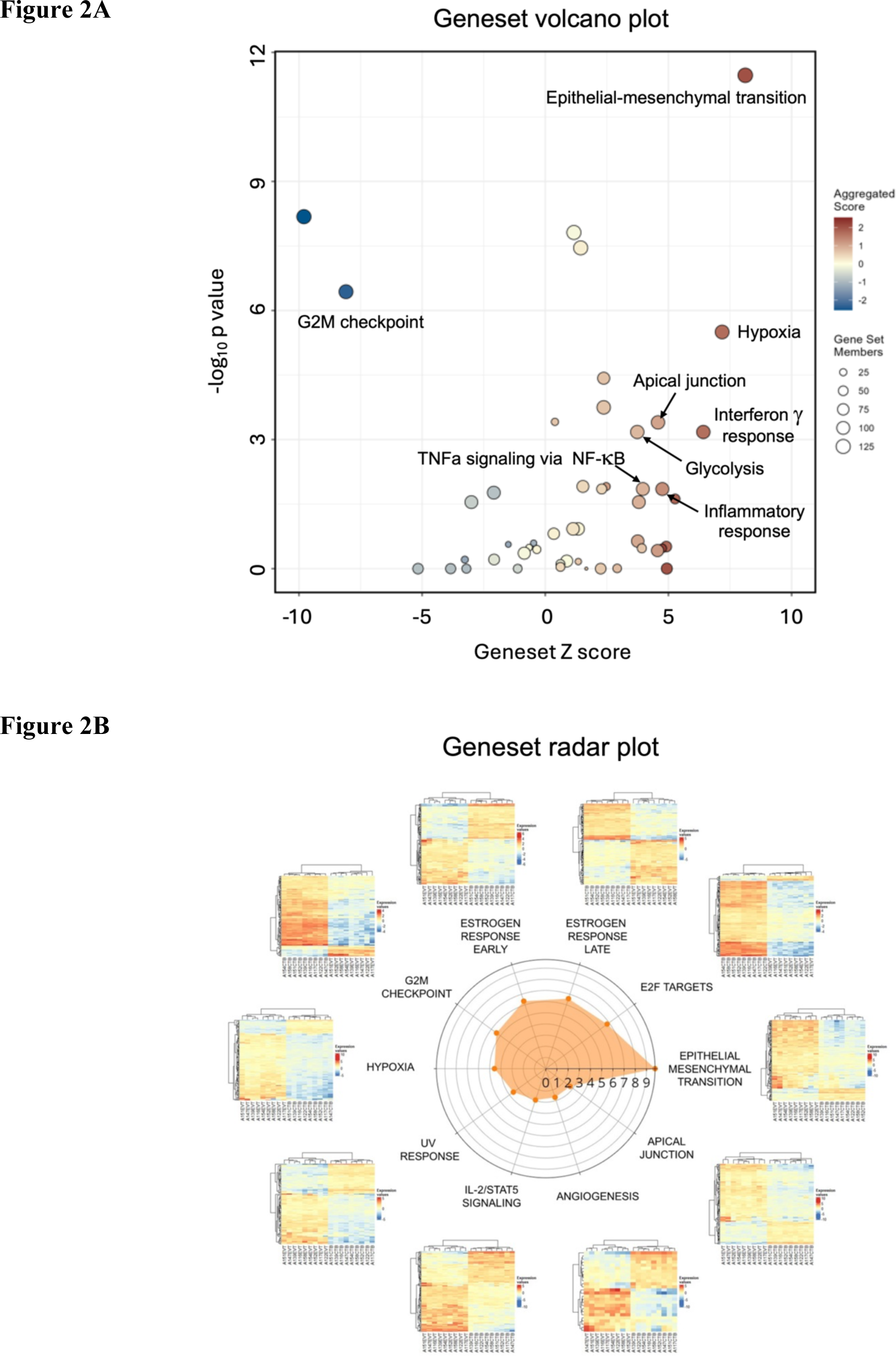
Functional enrichment of differentially expressed genes in EVT compared to CTB. ***(A)*** *Volcano/bubble plot of enrichment drawn from the Hallmark database of genesets**. (B)** Radar plot of the top 10 enriched genesets*.

**Table 1A.** Functional enrichment of biological processes in EVT compared to CTB.

| Biological process | Z score | Adj. p value |
| --- | --- | --- |
| HALLMARK_EPITHELIAL_MESENCHYMAL_TRANSITION | 8.12 | 1.71E-10 |
| HALLMARK_HYPOXIA | 7.18 | 2.63E-05 |
| HALLMARK_INTERFERON_GAMMA_RESPONSE | 6.42 | 2.77E-03 |
| HALLMARK_INFLAMMATORY_RESPONSE | 4.75 | 4.15E-02 |
| HALLMARK_APICAL_JUNCTION | 4.58 | 1.99E-03 |
| HALLMARK_TNFA_SIGNALING_VIA_NFKB | 3.96 | 4.15E-02 |
| HALLMARK_GLYCOLYSIS | 3.74 | 2.77E-03 |
| HALLMARK_TGF_BETA_SIGNALING | 2.47 | 4.15E-02 |
| HALLMARK_IL2_STAT5_SIGNALING | 2.37 | 1.11E-03 |
| HALLMARK_UV_RESPONSE_DN | 2.37 | 2.68E-04 |
| HALLMARK_ANDROGEN_RESPONSE | 2.29 | 4.15E-02 |
| HALLMARK_APOPTOSIS | 1.53 | 4.15E-02 |
| HALLMARK_ESTROGEN_RESPONSE_EARLY | 1.44 | 4.38E-07 |
| HALLMARK_MITOTIC_SPINDLE | -2.09 | 4.76E-02 |
| HALLMARK_G2M_CHECKPOINT | -8.09 | 3.66E-06 |

**Table 1B.** Enrichment of genes from the Hallmark EMT geneset in EVT compared to CTB.

| Symbol | Fold change | Symbol | Fold change | Symbol | Fold change |
| --- | --- | --- | --- | --- | --- |
| *SERPINE2* | 56.52 | *TNFAIP3* | 8.76 | *BMP1* | 3.80 |
| *QSOX1* | 48.45 | *COL12A1* | 8.64 | *ITGB1* | 3.70 |
| *COL16A1* | 37.57 | *PRRX1* | 8.34 | *MGP* | 3.61 |
| *COL11A1* | 36.29 | *PMP22* | 8.31 | *ABI3BP* | 3.54 |
| *FN1* | 35.59 | *PDLIM4* | 8.20 | *COL1A2* | 3.53 |
| *LRRC15* | 32.61 | *PLOD2* | 8.15 | *SAT1* | 3.53 |
| *PVR* | 31.10 | *PLOD3* | 8.07 | *PLOD1* | 3.53 |
| *HTRA1* | 30.31 | *NOTCH2* | 7.88 | *ECM1* | 3.49 |
| *ITGA5* | 29.39 | *IGFBP2* | 7.68 | *ITGA2* | 3.41 |
| *TIMP1* | 27.67 | *FSTL1* | 7.45 | *TFPI2* | 3.00 |
| *PMEPA1* | 26.96 | *CRLF1* | 7.11 | *PPIB* | 2.85 |
| *FSTL3* | 26.96 | *FBLN5* | 6.99 | *P3H1* | 2.65 |
| *SPARC* | 24.26 | *SDC4* | 6.94 | *TAGLN* | 2.55 |
| *MMP2* | 24.23 | *PCOLCE* | 6.88 | *SERPINH1* | 2.53 |
| *BGN* | 23.77 | *VCAN* | 6.80 | *FBN1* | 2.50 |
| *ADAM12* | 23.17 | *COL6A3* | 6.77 | *FBLN1* | 2.31 |
| *LAMA3* | 22.32 | *SLIT2* | 6.56 | *FAP* | 2.30 |
| *TPM1* | 22.30 | *TGFBI* | 6.35 | *MMP14* | 2.28 |
| *SLIT3* | 21.31 | *NNMT* | 6.18 | *CCN1* | 2.21 |
| *NID2* | 20.54 | *COL4A1* | 6.12 | *COPA* | 2.13 |
| *CAPG* | 20.46 | *APLP1* | 5.96 | *CD59* | 2.02 |
| *WIPF1* | 19.71 | *LUM* | 5.76 | *BASP1* | 1.99 |
| *GLIPR1* | 18.61 | *TNFRSF11B* | 5.70 | *FERMT2* | 1.87 |
| *LOXL1* | 18.16 | *DCN* | 5.65 | *CALD1* | 1.87 |
| *TGFB1* | 17.46 | *CDH11* | 5.64 | *LAMC1* | -1.75 |
| *CCN2* | 17.28 | *FBLN2* | 5.64 | *CAP2* | -2.04 |
| *TIMP3* | 16.01 | *COLGALT1* | 5.37 | *ITGAV* | -2.70 |
| *SERPINE1* | 15.38 | *IGFBP4* | 5.35 | *TGM2* | -3.13 |
| *INHBA* | 15.19 | *MATN2* | 5.17 | *JUN* | -3.33 |
| *NT5E* | 15.05 | *PDGFRB* | 5.05 | *BDNF* | -4.35 |
| *COL6A2* | 14.27 | *EMP3* | 4.89 | *COL5A2* | -4.76 |
| *LGALS1* | 13.66 | *CALU* | 4.68 | *LAMA1* | -5.00 |
| *IGFBP3* | 13.55 | *CXCL8* | 4.67 | *MCM7* | -5.56 |
| *RGS4* | 13.31 | *THY1* | 4.50 | *PCOLCE2* | -12.50 |
| *LOXL2* | 12.73 | *COL3A1* | 4.48 | *ID2* | -12.50 |
| *SFRP1* | 12.58 | *DKK1* | 4.39 | *FBN2* | -12.50 |
| *FLNA* | 11.97 | *PTHLH* | 4.38 | *DAB2* | -14.29 |
| *TPM2* | 11.19 | *COL4A2* | 4.37 | *ITGB5* | -20.00 |
| *EFEMP2* | 10.60 | *SPOCK1* | 4.29 | *MEST* | -33.33 |
| *MFAP5* | 10.06 | *MYL9* | 4.19 | *CADM1* | -100.00 |
| *COL5A1* | 9.85 | *VIM* | 4.01 |  |  |
| *PLAUR* | 9.73 | *SPP1* | 3.84 |  |  |

### Differences in EMT genes between basal plate and villous tree CTB

We investigated whether the changes seen in EVT compared to CTB might be a result of differences between the basal plate CTB, used here as controls for the DE analysis, and the villous tree CTB (vCTB). We find that while the PCA and clustergram (**Figure 1B**, **Figure 1D**) do not reveal obvious differences between CTB and vCTB, 459 genes show significant changes, 103 with increased expression in CTB, 355 showing a decrease. Some 22 genes overlapping with the EMT library show differential expression (**Supplementary Table S3**); 16 show an increase in CTB compared to vCTB and would be consistent with stimulation of EVT differentiation.

Importantly, these genes show further increases in EVT. Among these is the 5-fold increase in CTB compared to vCTB, of *ZEB1*, an EMT master regulator, which exhibits a further 5-fold increase in the paired EVT. Overall, these relatively limited changes indicate a slight EMT shift of the vCTB towards the mesenchymal end of the EMT spectrum.

### A subgroup of differentiation-associated genes have placenta-specific expression

There are a number of genes in our DE list which are primarily expressed in the placenta (Rawn and Cross, 2008) including, *CSH1*, *CSH2, PAPPA, GH2* and *HLA-G,* showing major increases in EVT compared to CTB (**Supplementary Table S4**). Also notable in that profile are substantial decreases in trophoblast genes expressed primarily in the syncytiotrophoblast (STB), including the retroviral genes which generate the syncytial fusion proteins syncytin-1 and syncytin-2 (*ERVW-1, ERVFRD-1*). Since EMT types 1, 2 and 3 have not been studied in trophoblast, the coordinated expression of these differentiation-associated genes with the DE- EMT genes (**Supplementary Table S2**) help to define a trophoblast differentiation signature.

For the purpose of rapid determination of trophoblast status, we have identified an abbreviated set of highly expressed genes including the EMT marker genes *FN1, ITGA5, MMP11, ADAM19* (upregulated in EVT), *BMP7, CTNNB1, MSX2, SERPINF1* (down-regulated in EVT), and differentiation-associated genes displaying similar characteristics including *CSH1* and *HLA-G* (up-regulated) and *ERVFRD-1* and *SIGLEC6* (down-regulated).

### Comparison of CTB and EVT methylomes highlights widespread losses and focal gains of CpG methylation in EVT

In parallel with the RNA-seq investigation, we analyzed DNA from EVT and CTB (n=4, 4) using Illumina Methylation EPIC Bead-Chip array (∼863K CpGs). Of the ∼23,000 genes covered in this assay, approximately 15,000 (∼190K CpGs on the Beadchip) demonstrated hypomethylation in the EVT compared to CTB, as indicated in the volcano plot (**Figure 3A**).

**Figure 3:**
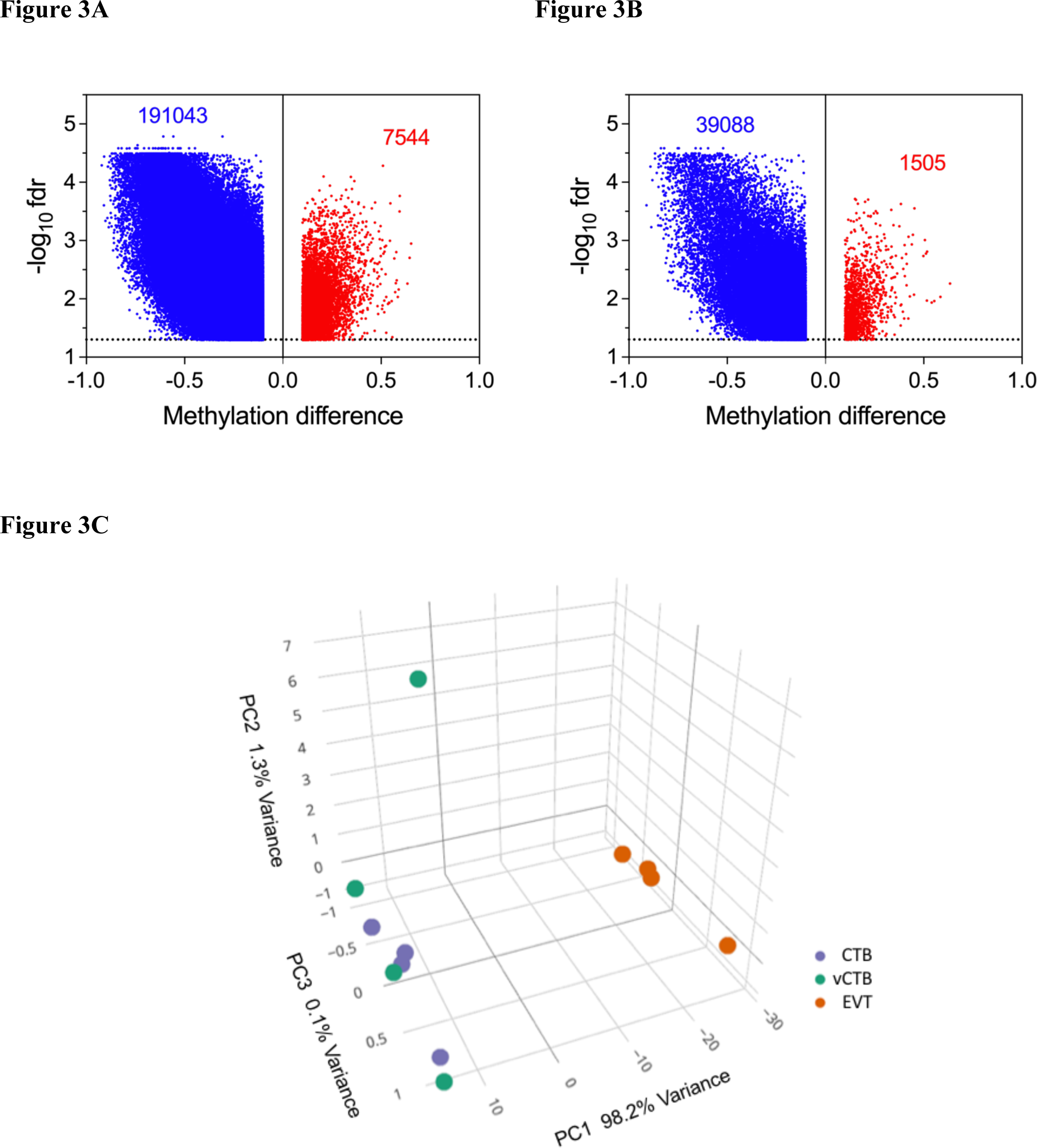
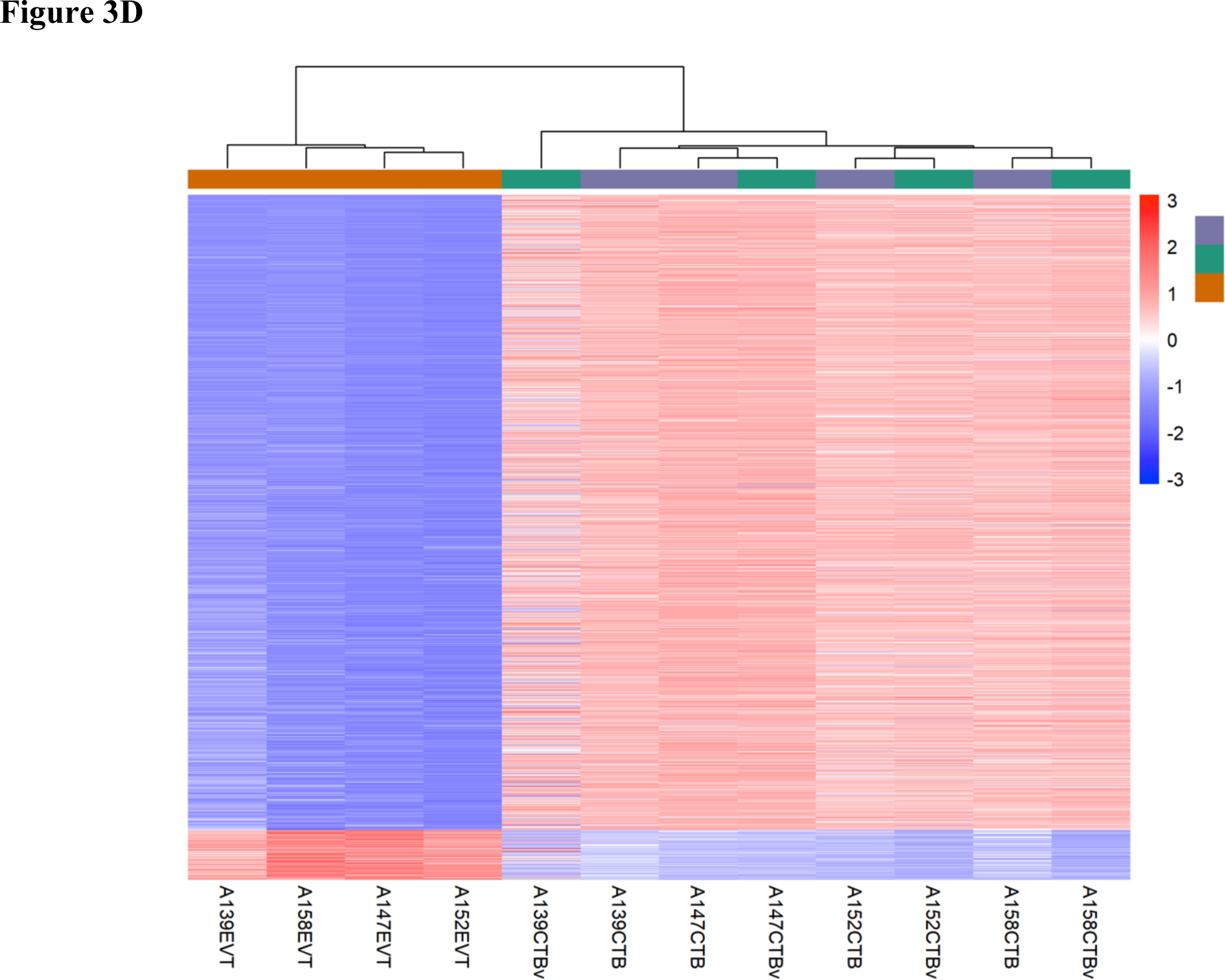
Differential methylation of genes in EVT compared to CTB. ***(A)*** *Volcano plot showing differences in total methylation (decreased – blue, increased – red) between CTB and EVT. **(B)** Volcano plot showing differences in promoter methylation (decreased – blue, increased – red) between CTB and EVT. **(C)** PCA plot of the methylome for CTB (purple), vCTB (green) and EVT (orange). **(D)** Clustergram showing differences in methylation between CTB, vCTB and EVT*

This shows the substantial asymmetry in the methylation changes between CTB and EVT, which is repeated in the subset of methylation changes taking place in gene promoter regions (**Figure 3B**). The principal components analysis of the methylation data (**Figure 3C**) exhibits separate clustering of CTB and EVT, similar to that shown for gene expression. There is an outlier in the CTB set however, a vCTB sample which is, nevertheless, well aligned with the other CTB samples in the PC1 dimension (accounting for 80% of the variance). Moreover, the clustergram for these samples (**Figure 3D**) shows clear differentiation, without any outliers, demonstrating separation between EVT and CTB. That hypomethylation dominates both cell types agrees with other studies showing the relative hypomethylation of the placenta or trophoblast compared to other somatic cells (Chatterjee et al., 2016; Ehrlich et al., 1982; Grigoriu et al., 2011; Novakovic et al., 2017; Schroeder et al., 2013; Zhang et al., 2021). Strikingly, our data show that the EVT are hypomethylated to an even greater degree than CTB. Our RNA-seq data shows that the *DNMT1*, *DNMT3A* and *DNMT3B* methyltransferase genes, responsible for much of DNA methylation, were down-regulated 7.7-, 4.2- and 3.8-fold respectively in the transition between CTB and EVT, perhaps explaining in part the broad hypomethylation observed in the EVT.

### Genes showing differential expression, methylation and involvement in the EMT

We also discovered a smaller set of genomic regions (1.6K CpGs on the Bead-Chips) demonstrating gain of methylation (GOM) in EVT relative to CTB. In contrast to the genome-wide hypomethylation described above, we suggest that these gains of methylation, which occur in regulatory sequences including promoter, enhancer, and insulator regions, might be specific, directed events. A gain of methylation (≥2 CpGs with ≥20 percent increase in methylation and a nominal FDR < 0.05) was associated with >1400 genes. Many of these GOM regions showed altered mRNA expression in the nearest genes. Meshing these GOM regions with our DE geneset revealed more than 700 genes that showed both gain of methylation and altered gene expression. Notably, the Hallmark EMT geneset was also enriched in this set of GOM-DE genes. The further intersection of these genes with those in the EMT library revealed a subset of 57 genes (GOM-DE-EMT genes; **Table 2**).

**Table 2:** Differentially expressed genes associated with EMT which show a gain of methylation.

| Gene ID | Fold change | CpG | Gene ID | Fold change | CpG | Gene ID | Fold change | CpG |
| --- | --- | --- | --- | --- | --- | --- | --- | --- |
| *MUC1* | 18.598 | 3 | *PLXND1* | 2.377 | 2 | *EML1* | -4.10 | 2 |
| *TGFB1* | 17.459 | 5 | *PKP3* | 1.739 | 4 | *IGF1R* | -4.20 | 4 |
| *FLT4* | 13.928 | 13 | *CSK* | 1.672 | 5 | *COL5A2* | -4.81 | 2 |
| *LOXL2* | 12.726 | 2 | *TGFBR3* | 1.651 | 2 | *ITGB4* | -4.90 | 5 |
| *MICALL2* | 12.545 | 16 |  |  |  | *BCL2* | -6.85 | 3 |
| *LPCAT1* | 10.175 | 2 | *BCL2L1* | -1.58 | 2 | *MET* | -7.75 | 2 |
| *SYNE1* | 9.949 | 3 | *SLC7A5* | -1.68 | 4 | *PLXNB1* | -8.62 | 2 |
| *COL5A1* | 9.855 | 78 | *KLF13* | -1.95 | 7 | *FXYD3* | -13.16 | 6 |
| *NFATC1* | 8.131 | 14 | *PTPN14* | -2.43 | 6 | *SPINT1* | -14.29 | 4 |
| *MBP* | 7.566 | 2 | *HRAS* | -2.60 | 5 | *ALDH1A3* | -14.71 | 4 |
| *RUNX1* | 7.538 | 38 | *MME* | -2.72 | 2 | *JAG1* | -15.63 | 2 |
| *FSCN1* | 6.221 | 9 | *TPD52L1* | -2.87 | 2 | *FGFR2* | -19.23 | 10 |
| *TEAD1* | 4.937 | 2 | *CTNNBIP1* | -2.97 | 3 | *ANK3* | -21.74 | 3 |
| *PRSS8* | 4.031 | 2 | *NOTCH1* | -2.99 | 13 | *BMP7* | -23.81 | 6 |
| *BMP1* | 3.796 | 2 | *MAP3K4* | -3.01 | 3 | *MSX2* | -47.62 | 2 |
| *FGFR3* | 3.708 | 20 | *TGM2* | -3.08 | 2 | *ELF5* | -125.00 | 4 |
| *TIAM1* | 3.698 | 2 | *CDK14* | -3.27 | 2 | *NRP2* | -125.00 | 3 |
| *AEBP1* | 3.451 | 3 | *VSNL1* | -3.40 | 2 | *SLC27A2* | -250.00 | 3 |
| *ABR* | 3.076 | 20 | *SYK* | -3.66 | 2 |  |  |  |
| *PKP2* | 3.029 | 2 | *EGFR* | -3.92 | 2 |  |  |  |

Among these genes were 24 where increased methylation was associated with increased gene expression and 33 where increased methylation was associated with decreased expression.

Examination of the biological processes associated with those GOM-DE-EMT genes showing increased methylation and gene expression, using the GOrilla gene ontology program, revealed that processes showing significant enrichment (FDR<0.05) were primarily concerned with promotion of an EMT and associated biological functions such as extracellular matrix organization (**Table 3**). By contrast, DE-GOM-DE-EMT genes which demonstrated decreased expression appeared to be clustered mainly around morphogenesis or epithelial development processes.

**Table 3:** Ontology for GOM-EMT genes with altered expression.

| Category | GO Term | Description | FDR<br>q-value | Enrichment |
| --- | --- | --- | --- | --- |
| Increased<br>expression | GO:1903055 | positive regulation of extracellular<br>matrix organization | 0.008 | 101.4 |
|  | GO:1903053 | regulation of extracellular matrix<br>organization | 0.002 | 66.1 |
|  | GO:0002062 | chondrocyte differentiation | 0.030 | 53.1 |
|  | GO:0055010 | ventricular cardiac muscle tissue<br>morphogenesis | 0.032 | 50.7 |
|  | GO:0010718 | positive regulation of epithelial to<br>mesenchymal transition | 0.033 | 44.6 |
|  | GO:0055008 | cardiac muscle tissue morphogenesis | 0.035 | 42.1 |
|  | GO:0034103 | regulation of tissue remodeling | 0.008 | 37.7 |
|  | GO:0060415 | muscle tissue morphogenesis | 0.045 | 36.6 |
|  | GO:0001837 | epithelial to mesenchymal transition | 0.050 | 33.8 |
|  | GO:0007043 | cell-cell junction assembly | 0.022 | 26.6 |
| Decreased<br>expression | GO:0060445 | branching involved in salivary gland<br>morphogenesis | 0.001 | 162.3 |
|  | GO:0002011 | morphogenesis of an epithelial sheet | 0.002 | 81.1 |
|  | GO:0033688 | regulation of osteoblast proliferation | 0.002 | 67.6 |
|  | GO:0031069 | hair follicle morphogenesis | 0.003 | 60.1 |
|  | GO:0014031 | mesenchymal cell development | 0.004 | 52.3 |
|  | GO:0110111 | negative regulation of animal organ<br>morphogenesis | 0.005 | 50.7 |
|  | GO:0090322 | regulation of superoxide metabolic<br>process | 0.005 | 46.4 |
|  | GO:0060411 | cardiac septum morphogenesis | 0.001 | 44.3 |
|  | GO:2000826 | regulation of heart morphogenesis | 0.006 | 43.9 |
|  | GO:0071526 | semaphorin-plexin signaling pathway | 0.006 | 43.9 |

### A 1.3 Mb domain gain of methylation in EVT spans the EMT-associated RUNX1 gene

In the examination of gain of methylation genes with increased gene expression, we observed a 1.3 Mb domain of extensive CpG hypermethylation in EVT, compared to CTB. This spanned the EMT-related *RUNX1* gene on human chromosome 21. Importantly, this broad domain of GOM was interrupted by smaller regions of loss of methylation, with the GOM being excluded from the major *RUNX1* promoter element, which was relatively *hypo*methylated in EVT (**Figure 4**). *RUNX1* shows a high gain of methylation during the CTB to EVT differentiation process, associated with a greater than 7-fold increase in mRNA expression (**Table 2)**. The increased methylation in *RUNX1* is largely accounted for by hypermethylation of the gene body (**Figure 4**), a process which has been shown to be positively correlated with increased transcription (Yang et al., 2014).

**Figure 4:**
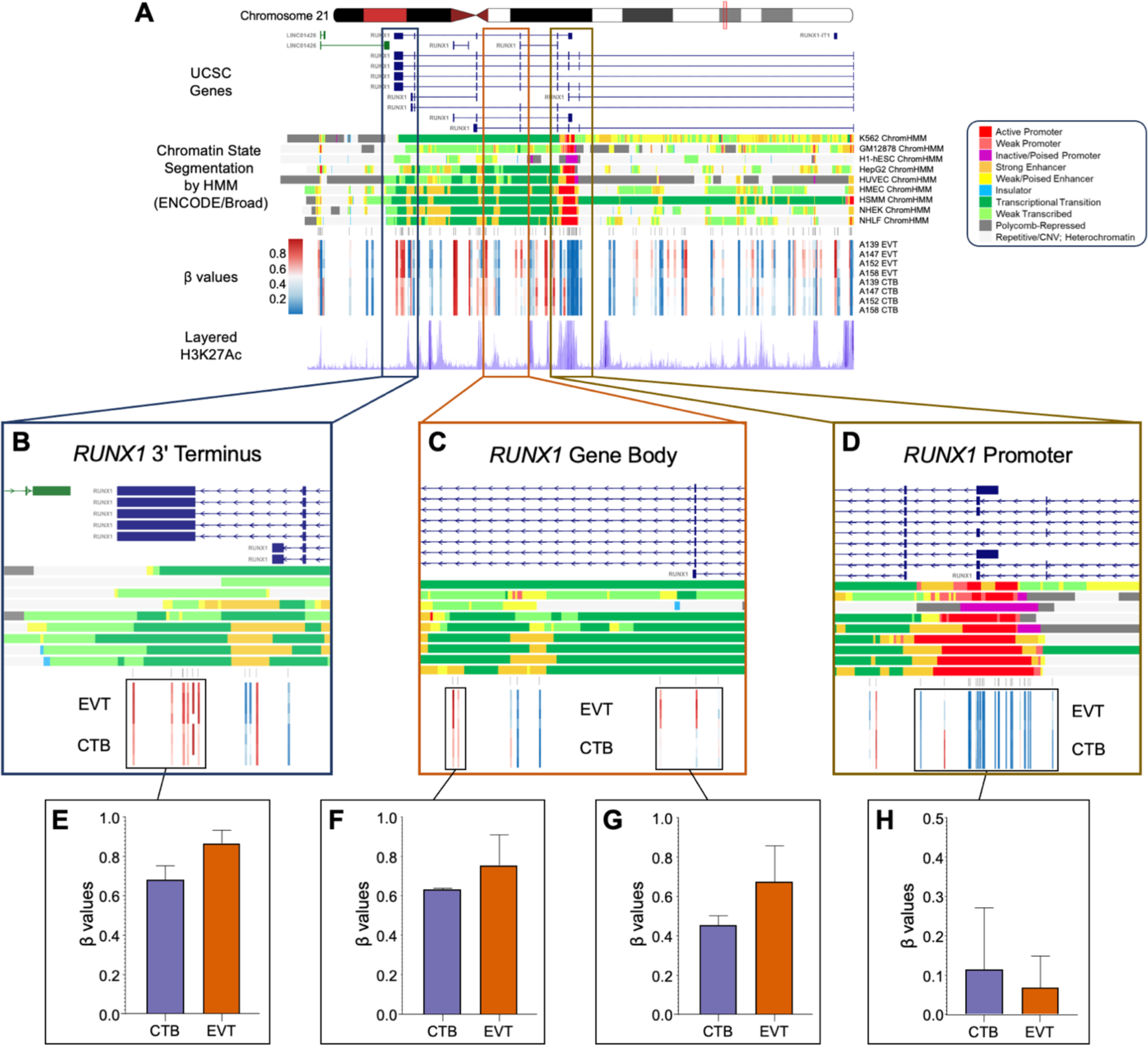
Regional RUNX1 gene body hypermethylation in EVTs relative to CTBs. ***(A)*** *Integrative genomics viewer (IGV) visualization of the genomic region around RUNX1 (Chr21: 36,110,847-36,448,197; hg19 coordinates) displaying the chromosome 21 ideogram and the following tracks: (1) UCSC known genes track showing RUNX1 transcripts (blue) and antisense LINC01426 transcripts (green); (2) Chromatin State Segmentation by Hidden Markov Model (ChromHMM) track for 9 ENCODE cell lines; (3) Illumina 850k EPIC methylation array track showing positions of CpG sites being measured; (4) heatmap representation of DNA methylation (β values) for EVT and CTB samples; (5) layered H3K27Ac track (epigenetic mark for active regulatory elements). **(B-D)** Enlarged views of three regions from the RUNX1 gene (3’ terminus, gene body, 5’ promoter) showing DNA methylation changes in relation to genomic and epigenomic features. (E-H) Bar graphs (mean ± SD) summarizing β values for select regions (boxes). The ChromHMM display conventions are as in* https://genome.ucsc.edu/cgi-bin/hgTrackUi?g=wgEncodeBroadHmm.

### Knock-down of RUNX1 leads to altered cell migration and invasion

One potential role for the GOM-DE-EMT genes is stabilizing an EVT phenotype leading to a restricted localization on the EMT spectrum. In third trimester EVT, this localization, between the epithelial and mesenchymal ends of the spectrum, appears to allow for maintenance of mesenchymal cell characteristics while abolishing cell invasiveness. This is supported by the activities described in the literature for non-trophoblastic cells, as shown in **Supplementary Table S5**, where we have listed the GOM-DE-EMT genes and their likely EMT roles.

To functionally test this concept, we investigated the role of *RUNX1* as an important transcription factor-encoding locus among the GOM-DE-EMT genes, which showed substantially increased methylation and a 7.5-fold increase in expression. We treated JEG3, a trophoblast cell line derived from an invasive choriocarcinoma, with an siRNA against *RUNX1* and investigated the effect on *RUNX1* gene expression, RUNX1 protein expression and JEG3 migratory capacity. As seen in **Figure 5A**, a 48 hr treatment of JEG3 with siRUNX1 reduced gene expression by ∼60% compared to treatment with the negative control (siNEG). **Figure 5B** shows a Western blot of siRUNX1-treated JEG3 cells using a monoclonal anti-RUNX1 antibody. The quantification of the Western blot is displayed in in **Figure 5C**, which shows an ∼55% reduction in RUNX1 protein in the siRUNX1-treated cells compared to siNEG-treated cells. After demonstrating the knockdown of *RUNX1* gene and RUNX1 protein, we measured the migratory capacity of siRUNX1-treated JEG3 cells using a scratch assay. **Figure 5D** shows that the migratory capacity of the siRUNX1-treated cells was reduced by ∼60% compared to the siNEG-treated controls at 72 hr., paralleling the knockdown in mRNA and protein. **Figure 5E** shows that knockdown of *RUNX1* mRNA and protein leads to reduced invasion through Matrigel, a different process than migration and consistent with the known role of *RUNX1* as a pro-EMT gene in other systems.

**Figure 5:**
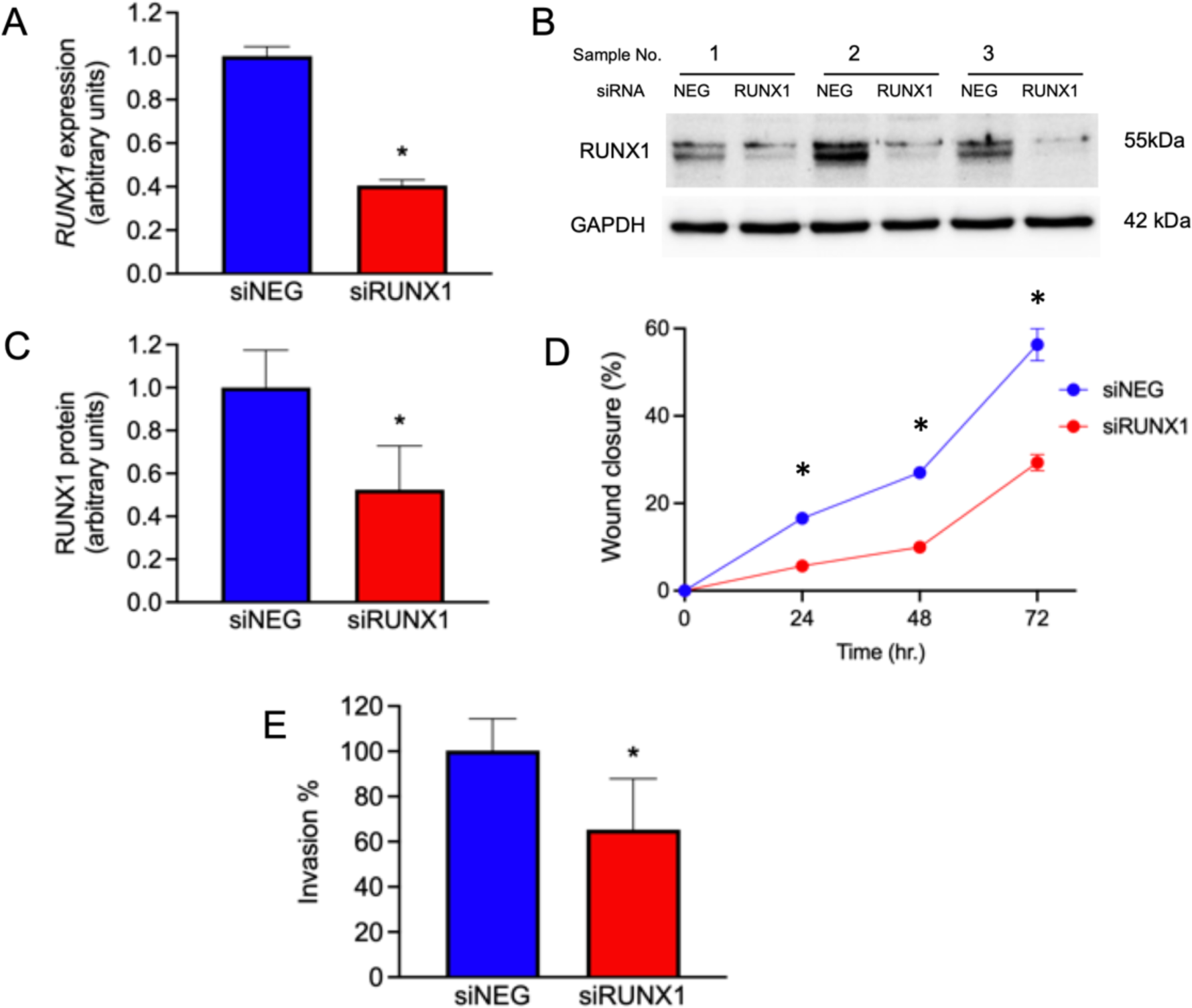
Role of RUNX1 in trophoblast. ***(A)*** *RUNX1 gene expression in JEG3 cells following treatment with siNEG or siRUNX1 (normalized to YWHAZ, n=4, * p < 0.05). **(B)** Western blot of JEG3 cells for RUNX1 and GAPDH following treatment with siNEG or siRUNX1. **(C)** Quantification of RUNX1 protein expression (normalized to GAPDH, n=3, * p < 0.05). **(D)** Cellular migration following treatment with siRUNX1, measured as a percentage of wound closure; n= 3; * p < 0.05. **(E)** Invasion of siNEG- or siRUNX1-treated JEG3 through Matrigel-coated Transwell membranes, measured as a percentage; n=16; * p < 0.001*

## Discussion

Our transcriptomic and methylomic comparison between term CTB and EVT has revealed substantial differences. In the transcriptomic comparison, we found over 7700 genes displaying gene expression changes. When this list was compared with a library of genes drawn from a broad set of EMT databases, there was >60% overlap This solidly supports the operation of an EMT mechanism in trophoblast differentiation. This is substantiated by the finding of the Hallmark EMT geneset as the most prominently enriched functional process. The EMT library did not contain identified trophoblast genes or those with trophoblast-specific promoters/regulators. These comprised another 26 genes which, when added to the broader EMT geneset, create a trophoblast-specific, third trimester EMT signature which, as we move to investigate invasion pathologies such as PAS and PE, will enable the assessment of EMT status in these conditions. Examining trophoblast methylation status revealed a substantial degree of hypomethylation in the EVT, even compared to the already-hypomethylated CTB. However, a small fraction of the sites tested demonstrated a gain of methylation. The intersection between these gain of methylation genes and the trophoblast EMT genes produced a list of EMT genes with both gain of methylation and differential expression. Functional testing of one of the primary genes in this group, *RUNX1*, confirmed its participation in the migration process characteristic of EVT, supporting that a small number of genes showing gain of methylation in EVT play a regulatory role in the trophoblast differentiation process.

### Confirmation of EMT in trophoblast differentiation

The substantial overlap of DE genes with the library of EMT genes (345 out of 563) demonstrates that an EMT is responsible, in large part, for the differentiation changes we observe. Our assessment aligns with the consensus guidelines from the EMT International Association (Yang et al., 2020). The well-described changes in cellular properties known to occur during the EMT and observed in CTB-to-EVT differentiation are associated with molecular changes observed with the EMT and stimulated by core transcription factors (DaSilva-Arnold et al., 2015; DaSilva-Arnold et al., 2019). Confirmation of an EMT raises the question of the location of third trimester EVT on the EMT spectrum. While first trimester EVT appear to be well-advanced towards the mesenchymal limit of the EMT spectrum (DaSilva-Arnold et al., 2015), third trimester EVT appear to have undergone regression back towards the epithelial pole (DaSilva-Arnold et al., 2018), a sign of the epithelial-mesenchymal plasticity which characterizes the EMT. Thus, the differential gene expression we observe in term placental EVT results of at least two major forces. The first is the initial drive towards differentiation which results in the mesenchymal phenotype of first trimester EVT. The second force is that which arrests and/or reverses the EVT progression, leading to the loss of invasiveness, loss of proliferative capacity and regression of EVT on the EMT spectrum. The data obtained here is, in all probability, an echo of the differentiation process which takes place early in gestation, blunted by the subsequent regression.

PCA derived from the DE genes (**Figure 2A**) clearly demonstrates the substantial difference between CTB and EVT. Significant also is the absence of major differences between vCTB and the CTB obtained from the placental basal plate, suggesting that the two have a similar phenotype. This is confirmed by the clustergram of the top 1000 genes shown in **Figure 2C** wherein the vCTB samples are intermixed with the CTB samples. This would suggest that vCTB and basal plate CTB are at a similar stage in the trophoblast lineage. Nevertheless, there are indications from EMT gene expression that basal plate CTB have, albeit in a small way, altered their expression profile toward the mesenchymal end of the EMT spectrum, as judged by the increases in *VIM* and the *ZEB1* master EMT regulator.

Examining the most significant genes driving the separation described by PC1 in the PCA plot (30 positive, 30 negative; **Figure 2B**), 17 are identified as part of the trophoblast EMT signature we have described. Of the other 43 genes, there are at least 21 which, while not part of the EMT databases, have been associated with the EMT in other contexts. The results derived from our PCA plot show great similarity to those obtained by Morey et al. (Morey et al., 2021), which show a comparable CTB/EVT separation in PC1 and a major overlap with the genes which drive this separation in our data. The predominance of EMT-associated genes driving the PC1 separation supports that the EMT is the primary process driving the differences between CTB and EVT.

### Changes in the DNA methylation profile

Several aspects of the changes in the DNA methylation profile between CTB and EVT stand out. The first is the overwhelming degree of hypomethylation in the EVT cells. The interesting comparison here is to the aberrant DNA methylation (i.e., genome-wide hypomethylation) observed in cancer cells (Nishiyama and Nakanishi, 2021). Within cancer cells, DNA methylation levels are reduced in regions of low CpG density compared with normal cells, while a subset of CpG islands are hypermethylated in a cell-specific manner, mainly targeting CpG islands in gene expression regulatory elements. This frequently results in alterations to genes controlling cell adhesion, migration and invasion (Nishiyama and Nakanishi, 2021), properties shared with EVT. The similarity to metastasizing cancer cells, which employ EMT as a transforming mechanism, again supports the role of the EMT in trophoblast differentiation.

The cause of the hypomethylation in the EVT is unknown. However, a possible contributing factor is reduced DNA methyltransferases, as both *DNMT3A* and *DNMT3B* show a gain of methylation within their promoter regions, while *DNMT1* shows a gain of methylation in other regulatory regions. All three methyltransferases show decreased gene expression in EVT. Our transcriptomic results also show a major decrease in *UHRF1*, which codes for a ubiquitination factor crucial for methylation. *UHRF1* depletion results in significant promoter demethylation and gene upregulation in cancer cells (Nishiyama and Nakanishi, 2021). However, this, like the actions of *TET1* and *TET2* which are normally associated with demethylation, requires the DNA replication-dependent dilution of DNA methylation marks by the inactivation of *de novo* DNA methylation. As third trimester CTB and EVT are largely non-proliferative, this indicates that demethylation processes observed for the EVT may occur earlier in gestation, at a time when these cells still retain proliferative capacity. These data would support the generation, early in gestation, of a mobile, invasive cell phenotype for the EVT, similar to cancer cell metastasis.

This is followed later by a loss of the invasive ability, which nevertheless occurs at a time when these cells are still proliferative, generating the hypomethylated EVT we observe in the third trimester. The timing of the loss of proliferative capacity is as yet unclear, but it marks the limit for DNA replication-dependent demethylation events. The methylation changes in CTB between early and late first trimester observed by Nordor et al (Nordor et al., 2017), may mark the point at which these transitions are occurring. This may also mark the point at which changes resulting in abnormal invasion occur.

Prior reports have addressed methylation in trophoblast differentiation. Most of these studies have shown a relative hypomethylation of placental tissue compared to other somatic cells (Chatterjee et al., 2016; Ehrlich et al., 1982; Fuke et al., 2004; Schroeder et al., 2013). These are confirmed by a more recent publications highlighting the relative hypomethylation of CTB relative to other tissues (Nordor et al., 2017; Zhang et al., 2021), although their gestational studies also indicate increasing CTB methylation moving from second to third trimester. There are also two reports which examined the extent of methylation separately in CTB and EVT (Gamage et al., 2018; Novakovic et al., 2017). The firstshows a small but significant difference between trophoblast and somatic cells, but a minimal difference between CTB and EVT. By contrast, the other study shows a clear reduction in EVT methylation compared to CTB, similar to the results here.

### Gain of methylation and trophoblast EMT

By contrast with the generalized hypomethylation, the EVT also show a gain of methylation in a small proportion of the sites surveyed, corresponding to over 1400 genes. The very limited extent in the gain of methylation, in the face of overwhelming EVT hypomethylation and the decreases in the expression of the methyltransferases, suggests to us that these events may be specifically directed, potentially with functional consequences. Within the DE geneset, a fraction show GOM and are identifiable as associated with the EMT (**Table 3**). An examination of these 57 genes reveals a mixture of pro-EMT and pro-MET (mesenchymal-epithelial transition) gene expression changes. For some genes, such as *TGFB1*, *LOXL2* and *RUNX1*, an increase in expression is associated with mechanisms promoting a pro-EMT shift, towards the mesenchymal phenotype (Cuevas et al., 2017; VanOudenhove et al., 2016), although there is also evidence, at least in breast cancer, that *RUNX1* may act as a tumor suppressor, a pro-MET role (Kulkarni et al., 2018). Others with increasing expression, such as *COL5A1* and *SYNE1*, are thought to be consequences of that shift rather than mediators, although they do form part of the mechano- transduction processes which accompany the EMT (Dejardin et al., 2020; Wang et al., 2022).

Most of the genes showing increased expression associated with a gain of methylation support a pro-EMT shift across the EMT spectrum (**Supplementary Table S5**). By contrast, for the methylated, EMT-associated genes showing a decrease in expression, the primary effects appear to support a pro-MET role. Overall, these two genesets demonstrate features which both promote and suppress the trophoblast EMT, an observation confirmed by the biological processes associated with the increased and decreased expression groups (**Table 3**).

### Functional effects of methylation

Despite the specific gains in methylation and an association with EMT, the consequences of these changes have not been tested functionally in trophoblast. Therefore, we chose to investigate the role of *RUNX1*, as an example of an EMT-associated gene demonstrating GOM and an increased DE in EVT vs. CTB. *Runx1* appears to play an important role in murine decidual development and the interaction of the decidua with invading trophoblast (Kannan et al., 2023). In the human placenta RUNX1 protein was observed in chorionic villi and decidua (Bermudez et al., 2021) but in the absence of co-labeling, cell type localization could not be confirmed. By contrast, the findings of Ponder et al point specifically to the presence of RUNX1 protein in cell column CTB and EVT throughout gestation (Ponder et al., 2016). *RUNX1* has also been shown to play a crucial role in the differentiation of hESC to early mesodermal lineages (VanOudenhove et al., 2016). In these latter experiments VanOudenhove et al discovered that depletion of *RUNX1* not only impaired the EMT, but also caused a loss in cellular motility, as reflected in a scratch assay. Our findings in JEG3 trophoblast lineage cells replicate this finding, and also demonstrate that loss of *RUNX1* mRNA and its encoded protein decreases trophoblast invasion, consistent with the general pro-EMT function of this gene. Our findings of increased *RUNX1* gene body methylation and upregulation of its mRNA in purified EVT, combined with the significant effects of its silencing on JEG3 cell migration, suggest that it plays an important role in trophoblast differentiation.

### Consequences of methylation and gene expression changes

We interpret these findings in the following manner. The EMT is controlled by the balance of forces driving cells towards the epithelial and mesenchymal ends of the EMT spectrum.

Changing the expression of genes controlling specific aspects of the EMT will force the cells towards one end or the other. Thus, in the first trimester there is high expression of genes such as *ZEB2* which promote transition towards the invasive, mesenchymal cell type (DaSilva-Arnold et al., 2015). As gestation progresses, other gene expression changes moderate this drive such that the cells become non-invasive but remain balanced between the more extreme epithelial and mesenchymal phenotypes.

We believe that the specific gains of methylation we observed may be part of a balancing act, preventing cells from regressing entirely to the epithelial phenotype while losing the invasive ability characteristic of the more mesenchymal, first trimester EVT. These gains in methylation would lock otherwise metastable cells into a non-proliferative, non-invasive phenotype.

Supporting this contention is data from Morey et al. (Morey et al., 2024) who show that the differences in methylation between primed and naïve iPSC-derived EVT appear to have significant consequences for invasiveness. What remains unknown at this point are those factors, acting through the (de)methylation pathways, which initiate and regulate the equilibrium of the third trimester EVT. The absence of decidual signals in tubal pregnancies, where invasion is not arrested, is perhaps instructive in this regard (Goffin et al., 2003; von Rango et al., 2003). The reduced number of multinuclear trophoblast giant cells, the fate of many EVT deeply embedded in the decidua/myometrium, in an over-invasive pathology such as accreta suggests that arrest and multinucleation may be linked, limiting mechanisms (Kemp et al., 2002; van Beekhuizen et al., 2009).

## Study limitations

The number of samples analyzed in this type of study is always a limitation, although we believe that the pairing of CTB and EVT derived from the same tissue may have compensated in part for that particular shortcoming. As with many studies of this type, the conclusions reached here are limited by the fact that we used only third trimester cells in this analysis. However, this study was designed to act as a baseline for third trimester investigations of pathological pregnancy conditions and has thus fulfilled its design expectations. Another limitation lies with the assessment of EMT genes. While we have used available libraries and databases, they are limited in their coverage and composed primarily of genes identified in cancer studies. There are however many genes which, while identified as being associated with the EMT in the literature, are not in our library.

The report by Morey et al. (Morey et al., 2021) has provided information on first and third trimester gene expression very similar to our third trimester data. Nevertheless, both have skirted another limitation which has become increasingly apparent from single cell RNA-seq studies, the possible presence of EVT cell subtypes or cell types regarded as intermediate between CTB and EVT extremes (Liu et al., 2018; Marsh et al., 2022; Tsang et al., 2017). The differences between vCTB and CTB provide some support for the existence of multiple subtypes, however the types, numbers and transience of these intermediates is a continuing issue.

## Conclusions

Our gene expression results confirm that the differentiation of CTB into EVT includes the substantial involvement of an EMT. We have generated a comprehensive, trophoblast-specific EMT signature. The differential DNA methylation profile demonstrates that the differentiation event is accompanied largely by EVT hypomethylation. The exception is the small number of genes showing a gain of methylation, half of which are associated with changes in gene expression. Within this latter group are genes which participate in the EMT, one example being *RUNX1*, which shows a gain of gene body CpG methylation. We demonstrated that *RUNX1* likely plays a significant role in the behavior of trophoblast cells, illustrated by changes in migratory rate and invasive efficiency upon *RUNX1* depletion in JEG3 cells. The GOM-DE- EMT genes show changes in expression which promote both EMT and its reverse, MET. This leads us to the concept that the change in the methylation profile is one means by which movement of these third trimester cells on the EMT spectrum is constrained, locking them into a stable configuration which is non-invasive but phenotypically mesenchymal.

## Methods

### Placental tissue and cell isolation

Placental tissue was obtained with written informed consent, as approved by the Hackensack University Medical Center IRB (Protocol Pro00002608). Tissue was obtained from normal, term (≥ 39 week’s gestational age) pregnancies, delivered by elective Caesarean section without labor. Tissue was transported to the laboratory, on ice, and processed within 20 minutes of delivery.

Excluded were pregnancies with medical or obstetric complications, mothers under 18 years of age or with a BMI ≥ 30.

EVT and CTB were isolated from these placentas as previously described (DaSilva-Arnold et al., 2018). A thin (3-4 mm) layer of tissue from the maternal-facing basal plate of the placenta was removed immediately following delivery. Tissue (∼20 g) was washed x 2 in phosphate-buffered saline (PBS), then further dissected into small pieces (2–4 mm) and washed x 2 in PBS. Following dissection, the placental tissue was incubated at 37°C in calcium- and magnesium-free Hank Balanced Salt Solution (HBSS) containing 10 mM Hepes, pH 7.4 with 0.05% trypsin (ThermoFisher Scientific, Waltham, MA), 0.1% dispase (Worthington Biochemical, Lakewood, NJ) and 12 U/mL Universal Nuclease (ThermoFisher). Tissue was incubated for 20 min in the digestion mix and then the supernatant was decanted, through a 100-μm filter, into FBS (final concentration 10%; Atlanta Biologicals, Flowery Branch GA) and placed on ice. Another aliquot of the digestion solution was added to the remaining tissue and the incubation and filtration steps were repeated x 2. Cells were pelleted by centrifugation (10 min, 1200 x g, 4°C) and resuspended in Separation Buffer (SB; calcium- and magnesium-free HBSS containing 2 mM EDTA, 0.5% BSA, pH 7.4 at 4°C). Lysis of red blood cells in the cell suspension was performed by rapid mixing of the cell suspension with an NH4Cl solution to obtain a final solution containing 155 mM NH4Cl, 10 mM KHCO3, 0.1 mM EDTA, pH 7.4. Following incubation of this lysis mixture for 10 min at room temperature, the cells were passed serially through 70 and 40 μm filters before centrifugation at 1200 x g for 10 min (4°C). Cells were resuspended in DMEM without calcium, containing 10% FBS, 1% penicillin/streptomycin and adjusted to pH 7.4 at 4°C (DMEM buffer). After a further wash in DMEM buffer, the cell mixture was divided into two aliquots which were incubated either with an anti- HLA-G antibody coupled to R-phycoerythrin (HLA-G-PE, 1:500, clone MEM G/9; Abcam, Waltham, MA) for isolation of

EVT, or with an anti-integrin ß4 antibody coupled to R-phycoerythrin (ITGß4-PE, 1:500, clone 58XB4, BioLegend, San Diego CA) for isolation of CTB. The cell mixture was incubated with antibody for 30 min at 4°C. Labeled cells were washed x 2 in DMEM buffer (400 x g, 5 min, 4°C), resuspended, counted and the volume was adjusted to 10^7^ cells/100 μL. Cells were incubated with anti-PE microbeads (10 μL microbeads/10^7^ cells; Miltenyi, San Diego, CA) for 20 min at 4°C, washed and resuspended in DMEM buffer. Isolation of HLA-G- and ITGß4- labeled cells was performed on an AutoMACS immunomagnetic cell separator (Miltenyi) using a double-column, positive selection procedure as specified by the manufacturer. In addition to the CTB isolated from the basal plate, CTB were also prepared from villous tissue, obtained from the area midway between placental basal and chorionic plates. These cells were prepared in the same way as the CTB isolated from the basal plate and are referred to henceforth as villous CTB (vCTB).

After separation, cells in the positive-selection fractions were counted, spun down (400 x g, 5 min, 4°C) and cell samples were taken to assess purity using flow cytometry. These samples were incubated with Sytox Green (30 nM, 20 min, room temperature; ThermoFisher) for assessment of cell death. Labeled cells were analyzed by flow cytometry, using a Cytomics FC500 flow cytometer (Beckman-Coulter, Indianapolis, IN). After analysis by flow cytometry, cells labeled by the anti-HLA-G antibody were found to comprise 95.6 ± 1.2% of the EVT fraction (n = 9). Cells labeled by the anti-integrin ß4 antibody comprised 96.8 ± 0.6% of the CTB fraction (n = 9). Cells were aliquoted for preparation of RNA (0.5 – 1.0 x 10^6^ cells/sample) or DNA (1.0 x 10^6^ cells/sample) and washed in PBS. Cells for preparation of RNA were pelleted then lysed immediately in Qiazol (Qiagen, Valencia, CA) and frozen at -80°C. Cells for DNA preparation were directly frozen at -80°C after pelleting.

### Cell culture and gene knockdown

JEG3 choriocarcinoma cells (ATCC, Manassas VA) were cultured at 37°C in DMEM/F12 containing 1% penicillin/streptomycin, 10% FBS (DMEM/F12). Cells (1 x 10^6^) were plated in each well of a 6-well plate and cultured to obtain a confluence of ∼70%. Cells were transfected with 30 nM siRNA (Silencer Select, ThermoFisher) directed against *RUNX1* (siRUNX1; #AM16708, assay ID 106564), or a negative control (siNEG; negative control #1), made up in Advanced DMEM/F12 (LDP, Cat# 12634010), and mixed with RNAiMAX (ThermoFisher) diluted in Advanced DMEMF/12 according to the manufacturer’s protocol. Following addition of the siRNA/RNAiMAX mixture, cells were incubated overnight in Advanced DMEM/F12 without antibiotics and then switched to complete DMEM/F12 for a total incubation period of 48 hr. At the end of the incubation period, cells were used in the migration/scratch assay and were also extracted, after washing, into Qiazol for qRT-PCR or into RIPA lysis buffer for Western blotting.

### Cell migration and invasion assays

For the Transwell invasion assays, Matrigel was thawed on ice and prepared at a concentration of 0.2 mg/mL, diluted in Advanced-DMEM/F12. Cold Fluoroblok inserts (VWR, Cat# 62406-504) were coated with 100 µL of Matrigel and incubated at 37°C overnight to allow the Matrigel to gel. Cell suspensions (∼60,000 cells per well; siNEG or siRUNX1 cells) were added to each of 4 wells. Subsequently, 0.5 mL of DMEM/F12 was added to the lower reservoir of each well and the plate was incubated for 48 hours. After incubation, calcein AM stock (Life Technologies, Cat# C1429; 1 mM in DMSO) was diluted to 2µM in warm HBSS/HEPES. The medium from the bottom reservoir was removed by aspiration, and the reservoir washed with HBSS/HEPES. Subsequently, 0.5 mL of the diluted calcein AM solution was added to the bottom reservoir of the wells and the plate was incubated at 37°C for 60 minutes. Plates were read from the bottom using a plate reader with an excitation wavelength of ∼488 nm and an emission wavelength of ∼520 nm.

The migration of JEG3 cells was measured using a scratch assay. After a 48 hr. transfection, JEG3 cells (siRUNX1 or siNEG ) were plated in a 12-well plate and grown to 70-80% confluence (18-24 hr.). Once at the requisite confluence, the cell layer was scraped in a straight line using a 1 mm sterile pipette tip. After scratching, the cell monolayer was washed to remove detached cells, then replenished with fresh medium. Wells were imaged by phase-contrast microscopy at 0, 24, 48 and 72 hr.

### RNA sequencing

RNA was extracted using the miRNeasy Micro kit (Qiagen) according to the manufacturer’s instructions. Genomic DNA was enzymatically digested by DNase I treatment and total RNA was captured by column purification. Both RNA concentration and integrity were quantified on an RNA 6000 Nano Chip using the Agilent 2100 Bioanalyzer (Agilent Technologies, Santa Clara, CA). These cell fractions, when extracted, yielded RNA with RIN scores of 8.7 ± 0.3 (EVT) and 9.4 ± 0.2 (CTB), displaying sufficient quality for RNA-sequencing analysis.

RNA sequencing was performed by the Genomics Center at Rutgers - New Jersey Medical School. Prior to RNA-sequencing, the quality of the RNA was checked using Agilent Tapestation. The RNA was ribosome-depleted using the Ribo-Zero kit (Illumina, San Diego, CA). Ribosome-depleted RNA was used to make cDNA libraries using NEBNext Ultra II RNA Library preparation kit for Illumina and NEBNext Multiplex Oligos for Illumina as per manufacturer’s protocol (New England Biolabs, Ipswich, MA). Briefly, ribosome-depleted RNA was chemically fragmented and reverse transcribed to make cDNA. The resulting cDNA was end-repaired and A-tailed. Adapters were ligated and libraries were PCR amplified using indexed primers. Libraries were then purified using Ampure XP beads (Beckman Coulter) and quantified using TapeStation and Qubit. Sequencing was performed on an Illumina NextSeq 500 with 1x75 configuration. The resulting raw data (bcl files) were converted to Fastq files and demultiplexed using Illumina software (bcl2fastq).

### RNA-sequencing data analysis

Quality control on raw Fastq files was performed using FastQC v0.11.8 (www.bioinformatics.babraham.ac.uk) and reads were mapped to the hg38 genome build with STAR v2.7.4a (Dobin et al., 2013) using the following parameters: --runMode alignReads -- outSAMtype BAM Unsorted --readFilesCommand zcat --genomeLoad LoadAndKeep -- clip3pAdapterSeq AGATCGGAAGAGC --clip3pAdapterMMp 1 –- outFilterScoreMinOverLread 0.3 –-outFilterMatchNminOverLread 0.3. Feature counts were generated using SAMtools v1.10 (Danecek et al., 2021) in combination with HTSeq v0.11.2 (Anders et al., 2015); specifically, the output of the “samtools view” command was piped to the “htseq-count” command with the –stranded=reverse parameter in the Python v3.6.1 environment. Human Genome Organisation Gene Nomenclature Committee (HGNC) symbols were updated using the R package HGNChelper v0.8.1 (Oh et al., 2020).

### Differential Abundance Analysis

Statistical testing for differential transcript abundance was performed with DESeq2 v1.34.0 (Love et al., 2014) in R using models that included the cell type and subject identifier (placenta of origin). Assigned fetal sex was considered separately since inclusion together with subject information provided a design formula that was not full rank, such that models could not be fit. Statistical significance was considered to be a false discovery rate (FDR) < 0.05 using the Benjamini-Hochberg procedure.

### Functional Enrichment Analysis and Visualization

Functional enrichment was assessed by overrepresentation analysis (hypergeometric test) using the enricher() function in the clusterProfiler v4.2.2 R package (Yu et al., 2012). Specifically, we tested for enrichment of the Hallmark gene sets (v7.4) obtained from the Molecular Signatures Database (MSigDB) collections using differentially abundant transcripts (FDR<0.05) compared to the background of all mapped transcripts; parameters included gene sets between 10 and 250 members in size, a threshold p-value of 0.05, and the Benjamini-Hochberg procedure for p-value adjustment.

The GeneTonic v1.6.1 workflow (Marini et al., 2021) was used for interactive visualization of the enrichment results. In this environment, the expression and statistical results tables generated by DESeq2 were combined with the enrichment analysis results obtained from clusterProfiler and an annotation data table acquired using the AnnotationDbi v1.56.2 package and the Bioconductor annotation package org.Hs.eg.db v3.14.0. Gene-geneset bipartite network graphs were produced using the ggs_graph() function and visualized using the visIgraph() function in the visNetwork library (https://CRAN.R-project.org/package=visNetwork). Individual heatmaps for enriched hallmark pathways were generated using the gs_heatmap() function. Radar plots and volcano plots were created with the gs_radar() and gs_volcano() functions, respectively.

Alternate renderings of these results were generated with Prism v.9.3 (GraphPad Software, San Diego, CA).

Biological process enrichment analysis was also performed using GOrilla (***G****ene **O**ntology en**RI**chment ana**L**ysis and visua**L**iz**A**tion tool;* (Eden et al., 2007; Eden et al., 2009)). A list of target genes was utilized in conjunction with a background list comprising all mapped transcripts.

### DNA Methylation

Analysis of DNA methylation was performed as described previously (Mendioroz et al., 2015), using Illumina Infinium Methylation EPIC v1 BeadChips, High molecular weight DNA was prepared from the frozen cell pellets by standard SDS/proteinase-K lysis followed by precipitation in 80 % isopropanol with glycogen as a carrier. Genomic DNA quality was evaluated by gel electrophoresis and quantity was determined using the PicoGreen DNA quantification assays (Life Technologies). Analysis of the DNA was performed using Illumina Infinium Methylation EPIC v1 BeadChips, according to the manufacturer’s instructions. The methylation assays were carried out at the Roswell Park Cancer Institute Genomics Shared Resource. BeadChip-based methylation assays involve bisulfite conversion of the genomic DNA, in which unmethylated cytosines are converted to uracil. This is followed by primer extensions to query the percentage methylation at each of 862,927 (850K) CpG dinucleotides, covering CpG islands (CGIs) within and around both promoter and non-promoter regions as well as many non-island promoter regions, associated with 99% of RefSeq genes. The resulting dataset was processed with GenomeStudio (v2011.1) software using the Methylation Module (v1.9.0). After background correction and normalization to internal controls, the percentage methylation (AVG_Beta) at each CpG queried by the array was calculated. Poorly performing probes with missing values (AVG_Beta detection p > 0.05 occurring in more than one sample per subgroup) and probes mapping to the X or Y chromosome were removed.

### qRT-PCR

*RUNX1* mRNA was measured by qRT-PCR. cDNA was synthesized from 200 ng of RNA using the High-Capacity cDNA Reverse Transcription Kit (Cat No. 4368814; ThermoFisher). qPCR was performed using primers for *RUNX1* (Hs01021970_m1; ThermoFisher) and the *YWHAZ* housekeeping gene (Hs03044281_g1) and the TaqMan™ Fast Advanced Master Mix for qPCR (Cat. No.: 4444556; ThermoFisher) on a QuantStudio 6 Flex Real-Time PCR unit (ThermoFisher). Changes in *RUNX1* expression were calculated using the 2-^ΔΔCT^ methodology.

### Western blotting

After washing with PBS, cells were extracted into RIPA lysis buffer. Protein (30 µg/well) was loaded on to 8% SDS gels for electrophoresis. Proteins were transferred from the gels to a nitrocellulose membrane (Bio-Rad, , Hercules, CA), blocked (0.5% fat dried milk in PBST, 60 min) and incubated with an anti-RUNX1 polyclonal antibody (1:1000; GTX129100, Genetex, Irvine, CA) or anti-ß-actin polyclonal antibody (1:1000; Cat# 4967S, Cell Signaling Technology, Danvers, MA) at 4°C overnight followed, after washing x 3 with PBST, by a 60 min incubation with an anti-rabbit IgG coupled to HRP (1:1000; Cat# 4967S, Cell Signaling Technology). Blots were incubated with Pierce ECL Western Blotting Substrate (ThermoFisher) and visualized using a ChemiDoc MP imager (Bio-Rad). Blots were quantitated using ImageJ (v1.64, NIH).

### Statistical analysis

Sex- and gestational-age specific birthweight centiles were calculated according to Fenton and Kim (Fenton and Kim, 2013). Data from the qRT-PCR assays, Western blots and scratch assays was checked for normality using the Shapiro-Wilk test. Statistical analysis was carried out using either an unpaired, two-tailed *t* test or a two-tailed Mann-Whitney U test, as appropriate, using Graphpad Prism v9.3.

## Acknowledgements

The authors would like to acknowledge the faculty and staff of the Department of Obstetrics and Gynecology at Hackensack University Medical Center for their assistance in obtaining the placental tissue samples used in this study.

## Contributions

Conceptualization (NI, SZ, SD, BT), methodology (NI, BT, SD, WA), software (WA, MR, CD), validation (SD, WA, MR), formal analysis (WA, MR, CD ), investigation (NI, SD, MT, MS, AC), resources (SD, SZ), data curation (WA, MR), writing - original draft preparation (NI), writing – review and editing (NI, BT, WA, SZ, SD), visualization (NI, WA, BT), supervision (NI, BT).

## Data Availability

The RNA-seq and DNA methylation data can be found in GEO under the accession numbers GSE256412 and GSE259304 respectively

## References

1. Anders, S., Pyl, P. T. and Huber, W. (2015). HTSeq--a Python framework to work with high- throughput sequencing data. Bioinformatics 31, 166–169.10.1093/bioinformatics/btu638 PMC4287950

2. Anton, L., Brown, A. G., Bartolomei, M. S. and Elovitz, M. A. (2014). Differential methylation of genes associated with cell adhesion in preeclamptic placentas. PLoS ONE [Electronic Resource] 9, e100148

3. Bermudez, L. G., Madariaga, I., Zuniga, M. I., Olaya, M., Canas, A., Rodriguez, L. S., Moreno, O. M. and Rojas, A. (2021). RUNX1 gene expression changes in the placentas of women smokers. Exp Ther Med 22, 902.10.3892/etm.2021.10334 PMC8243315

4. Chatterjee, A., Macaulay, E. C., Rodger, E. J., Stockwell, P. A., Parry, M. F., Roberts, H. E., Slatter, T. L., Hung, N. A., Devenish, C. J. and Morison, I. M. (2016). Placental Hypomethylation Is More Pronounced in Genomic Loci Devoid of Retroelements. G3(Bethesda) 6, 1911-1921.10.1534/g3.116.030379 PMC4938645

5. Chen, Y., Wang, K., Qian, C. N. and Leach, R. (2013). DNA methylation is associated with transcription of Snail and Slug genes. Biochem Biophys Res Commun 430, 1083–1090.10.1016/j.bbrc.2012.12.034 PMC3552036

6. Cheng, W. Y., Kandel, J. J., Yamashiro, D. J., Canoll, P. and Anastassiou, D. (2012). A multi-cancer mesenchymal transition gene expression signature is associated with prolonged time to recurrence in glioblastoma. PLoS One 7, e34705.10.1371/journal.pone.0034705 PMC3321034

7. Cuevas, E. P., Eraso, P., Mazon, M. J., Santos, V., Moreno-Bueno, G., Cano, A. and Portillo, F. (2017). LOXL2 drives epithelial-mesenchymal transition via activation of IRE1-XBP1 signalling pathway. Sci Rep 7, 44988.10.1038/srep44988 PMC5362953

8. Danecek, P., Bonfield, J. K., Liddle, J., Marshall, J., Ohan, V., Pollard, M. O., Whitwham, A., Keane, T., McCarthy, S. A., Davies, R. M. and Li, H. (2021). Twelve years of SAMtools and BCFtools. Gigascience 10.10.1093/gigascience/giab008 PMC7931819

9. DaSilva-Arnold, S., James, J. L., Al-Khan, A., Zamudio, S. and Illsley, N. P. (2015). Differentiation of first trimester cytotrophoblast to extravillous trophoblast involves an epithelial-mesenchymal transition. Placenta 36, 1412–1418.10.1016/j.placenta.2015.10.013

10. DaSilva-Arnold, S., Zamudio, S., Al-Khan, A., Alvarez-Perez, J., Mannion, C., Koenig, C., Luke, D., Perez, A., Petroff, M., Alvarez, M. and Illsley, N. P. (2018). Human trophoblast epithelial-mesenchymal transition in abnormally invasive placenta. Biol Reprod 99, 409–421.10.1093/biolre/ioy042

11. DaSilva-Arnold, S. C., Kuo, C. Y., Davra, V., Remache, Y., Kim, P. C. W., Fisher, J. P., Zamudio, S., Al-Khan, A., Birge, R. B. and Illsley, N. P. (2019). ZEB2, a master regulator of the epithelial-mesenchymal transition, mediates trophoblast differentiation. Mol Hum Reprod 25, 61–75.10.1093/molehr/gay053 PMC6497037

12. Davies, J. E., Pollheimer, J., Yong, H. E., Kokkinos, M. I., Kalionis, B., Knofler, M. and Murthi, P. (2016). Epithelial-mesenchymal transition during extravillous trophoblast differentiation. Cell Adh Migr 10, 310–321.10.1080/19336918.2016.1170258 PMC4951171

13. Dejardin, T., Carollo, P. S., Sipieter, F., Davidson, P. M., Seiler, C., Cuvelier, D., Cadot, B., Sykes, C., Gomes, E. R. and Borghi, N. (2020). Nesprins are mechanotransducers that discriminate epithelial-mesenchymal transition programs. J Cell Biol 219.10.1083/jcb.201908036 PMC7659719

14. Dobin, A., Davis, C. A., Schlesinger, F., Drenkow, J., Zaleski, C., Jha, S., Batut, P., Chaisson, M. and Gingeras, T. R. (2013). STAR: ultrafast universal RNA-seq aligner. Bioinformatics 29, 15–21.10.1093/bioinformatics/bts635 PMC3530905

15. Eden, E., Lipson, D., Yogev, S. and Yakhini, Z. (2007). Discovering motifs in ranked lists of DNA sequences. PLoS Comput Biol 3, e39.10.1371/journal.pcbi.0030039 PMC1829477

16. Eden, E., Navon, R., Steinfeld, I., Lipson, D. and Yakhini, Z. (2009). GOrilla: a tool for discovery and visualization of enriched GO terms in ranked gene lists. BMC Bioinformatics 10, 48.10.1186/1471-2105-10-48 PMC2644678

17. Ehrlich, M., Gama-Sosa, M. A., Huang, L. H., Midgett, R. M., Kuo, K. C., McCune, R. A. and Gehrke, C. (1982). Amount and distribution of 5-methylcytosine in human DNA from different types of tissues of cells. Nucleic Acids Res 10, 2709–2721.10.1093/nar/10.8.2709 PMC320645

18. Fenton, T. R. and Kim, J. H. (2013). A systematic review and meta-analysis to revise the Fenton growth chart for preterm infants. BMC Pediatr 13, 59.10.1186/1471-2431-13-59 PMC3637477

19. Fuke, C., Shimabukuro, M., Petronis, A., Sugimoto, J., Oda, T., Miura, K., Miyazaki, T., Ogura, C., Okazaki, Y. and Jinno, Y. (2004). Age related changes in 5-methylcytosine content in human peripheral leukocytes and placentas: an HPLC-based study. Ann Hum Genet 68, 196–204.10.1046/j.1529-8817.2004.00081.x

20. Gamage, T., Schierding, W., Tsai, P., Ludgate, J. L., Chamley, L. W., Weeks, R. J., Macaulay, E. C. and James, J. L. (2018). Human trophoblasts are primarily distinguished from somatic cells by differences in the pattern rather than the degree of global CpG methylation. Biol Open 7.10.1242/bio.034884 PMC6124577

21. Goffin, F., Munaut, C., Malassine, A., Evain-Brion, D., Frankenne, F., Fridman, V., Dubois, M., Uzan, S., Merviel, P. and Foidart, J. M. (2003). Evidence of a limited contribution of feto-maternal interactions to trophoblast differentiation along the invasive pathway. Tissue Antigens 62, 104–116

22. Grigoriu, A., Ferreira, J. C., Choufani, S., Baczyk, D., Kingdom, J. and Weksberg, R. (2011). Cell specific patterns of methylation in the human placenta. Epigenetics 6, 368–379.10.4161/epi.6.3.14196 PMC3092685

23. Groger, C. J., Grubinger, M., Waldhor, T., Vierlinger, K. and Mikulits, W. (2012). Meta- analysis of gene expression signatures defining the epithelial to mesenchymal transition during cancer progression. PLoS One 7, e51136.10.1371/journal.pone.0051136 PMC3519484

24. Illsley, N. P., DaSilva-Arnold, S. C., Zamudio, S., Alvarez, M. and Al-Khan, A. (2020). Trophoblast invasion: Lessons from abnormally invasive placenta (placenta accreta). Placenta 102, 61–66.10.1016/j.placenta.2020.01.004 PMC7680503

25. Kalluri, R. and Weinberg, R. A. (2009). The basics of epithelial-mesenchymal transition. J Clin Invest 119, 1420–1428.10.1172/JCI391042689101

26. Kannan, A., Beal, J. R., Neff, A. M., Bagchi, M. K. and Bagchi, I. C. (2023). Runx1 regulates critical factors that control uterine angiogenesis and trophoblast differentiation during placental development. PNAS Nexus 2, pgad215.10.1093/pnasnexus/pgad215 PMC10321400

27. Kemp, B., Kertschanska, S., Kadyrov, M., Rath, W., Kaufmann, P. and Huppertz, B. (2002). Invasive depth of extravillous trophoblast correlates with cellular phenotype: a comparison of intra- and extrauterine implantation sites. Histochem Cell Biol 117, 401–414.10.1007/s00418-002-0396-0

28. Knofler, M., Haider, S., Saleh, L., Pollheimer, J., Gamage, T. and James, J. (2019). Human placenta and trophoblast development: key molecular mechanisms and model systems. Cell Mol Life Sci 76, 3479–3496.10.1007/s00018-019-03104-6 PMC6697717

29. Kokkinos, M. I., Murthi, P., Wafai, R., Thompson, E. W. and Newgreen, D. F. (2010). Cadherins in the human placenta--epithelial-mesenchymal transition (EMT) and placental development. Placenta 31, 747–755.10.1016/j.placenta.2010.06.017

30. Kulkarni, M., Tan, T. Z., Syed Sulaiman, N. B., Lamar, J. M., Bansal, P., Cui, J., Qiao, Y. and Ito, Y. (2018). RUNX1 and RUNX3 protect against YAP-mediated EMT, stem-ness and shorter survival outcomes in breast cancer. Oncotarget 9, 14175–14192.10.18632/oncotarget.24419 PMC5865662

31. Liu, Y., Fan, X., Wang, R., Lu, X., Dang, Y. L., Wang, H., Lin, H. Y., Zhu, C., Ge, H., Cross, J. C. and Wang, H. (2018). Single-cell RNA-seq reveals the diversity of trophoblast subtypes and patterns of differentiation in the human placenta. Cell Res 28, 819–832.10.1038/s41422-018-0066-y PMC6082907

32. Love, M. I., Huber, W. and Anders, S. (2014). Moderated estimation of fold change and dispersion for RNA-seq data with DESeq2. Genome Biol 15, 550.10.1186/s13059-014- 0550-8 PMC4302049

33. Marini, F., Ludt, A., Linke, J. and Strauch, K. (2021). GeneTonic: an R/Bioconductor package for streamlining the interpretation of RNA-seq data. BMC Bioinformatics 22, 610.10.1186/s12859-021-04461-5 PMC8697502

34. Marsh, B., Zhou, Y., Kapidzic, M., Fisher, S. and Blelloch, R. (2022). Regionally distinct trophoblast regulate barrier function and invasion in the human placenta. Elife 11.10.7554/eLife.78829 PMC9323019

35. Mendioroz, M., Do, C., Jiang, X., Liu, C., Darbary, H. K., Lang, C. F., Lin, J., Thomas, A., Abu-Amero, S., Stanier, P., et al. (2015). Trans effects of chromosome aneuploidies on DNA methylation patterns in human Down syndrome and mouse models. Genome Biol 16, 263.10.1186/s13059-015-0827-6 PMC4659173

36. Morey, R., Bui, T., Cheung, V. C., Dong, C., Zemke, J. E., Requena, D., Arora, H., Jackson, M. G., Pizzo, D., Theunissen, T. W. and Horii, M. (2024). iPSC-based modeling of preeclampsia identifies epigenetic defects in extravillous trophoblast differentiation. iScience 27, 109569.10.1016/j.isci.2024.109569 PMC11016801

37. Morey, R., Farah, O., Kallol, S., Requena, D. F., Meads, M., Moretto-Zita, M., Soncin, F., Laurent, L. C. and Parast, M. M. (2021). Transcriptomic Drivers of Differentiation, Maturation, and Polyploidy in Human Extravillous Trophoblast. Front Cell Dev Biol 9, 702046.10.3389/fcell.2021.702046 PMC8446284

38. Natenzon, A., McFadden, P., DaSilva-Arnold, S. C., Zamudio, S. and Illsley, N. P. (2022). Diminished trophoblast differentiation in early onset preeclampsia. Placenta 120, 25–31.10.1016/j.placenta.2022.02.004

39. Nishiyama, A. and Nakanishi, M. (2021). Navigating the DNA methylation landscape of cancer. Trends Genet 37, 1012–1027.10.1016/j.tig.2021.05.002

40. Nordor, A. V., Nehar-Belaid, D., Richon, S., Klatzmann, D., Bellet, D., Dangles-Marie, V., Fournier, T. and Aryee, M. J. (2017). The early pregnancy placenta foreshadows DNA methylation alterations of solid tumors. Epigenetics 12, 793–803.10.1080/15592294.2017.1342912 PMC5739102

41. Novakovic, B., Evain-Brion, D., Murthi, P., Fournier, T. and Saffery, R. (2017). Variable DAXX gene methylation is a common feature of placental trophoblast differentiation, preeclampsia, and response to hypoxia. FASEB J 31, 2380–2392.10.1096/fj.201601189RR

42. Oh, S., Abdelnabi, J., Al-Dulaimi, R., Aggarwal, A., Ramos, M., Davis, S., Riester, M. and Waldron, L. (2020). HGNChelper: identification and correction of invalid gene symbols for human and mouse. F1000Res 9, 1493.10.12688/f1000research.28033.2 PMC7856679

43. Phipps, E. A., Thadhani, R., Benzing, T. and Karumanchi, S. A. (2019). Pre-eclampsia: pathogenesis, novel diagnostics and therapies. Nature Reviews Nephrology 15, 275–289.10.1038/s41581-019-0119-6

44. Ponder, K. L., Barcena, A., Bos, F. L., Gormley, M., Zhou, Y., Ona, K., Kapidzic, M., Zovein, A. C. and Fisher, S. J. (2016). Preeclampsia and Inflammatory Preterm Labor Alter the Human Placental Hematopoietic Niche. Reprod Sci 23, 1179–1192.10.1177/1933719116632926 PMC5933163

45. Rawn, S. M. and Cross, J. C. (2008). The evolution, regulation, and function of placenta- specific genes. Annu Rev Cell Dev Biol 24, 159–181.10.1146/annurev.cellbio.24.110707.175418

46. Schroeder, D. I., Blair, J. D., Lott, P., Yu, H. O., Hong, D., Crary, F., Ashwood, P., Walker, C., Korf, I., Robinson, W. P. and LaSalle, J. M. (2013). The human placenta methylome. Proc Natl Acad Sci U S A 110, 6037–6042.10.1073/pnas.1215145110 PMC3625261

47. Tantbirojn, P., Crum, C. P. and Parast, M. M. (2008). Pathophysiology of placenta creta: the role of decidua and extravillous trophoblast. Placenta 29, 639–645.10.1016/j.placenta.2008.04.008

48. Taube, J. H., Herschkowitz, J. I., Komurov, K., Zhou, A. Y., Gupta, S., Yang, J., Hartwell, K., Onder, T. T., Gupta, P. B., Evans, K. W., et al. (2010). Core epithelial-to- mesenchymal transition interactome gene-expression signature is associated with claudin- low and metaplastic breast cancer subtypes. Proc Natl Acad Sci U S A 107, 15449–15454.10.1073/pnas.1004900107 PMC2932589

49. Tsang, J. C. H., Vong, J. S. L., Ji, L., Poon, L. C. Y., Jiang, P., Lui, K. O., Ni, Y. B., To, K. F., Cheng, Y. K. Y., Chiu, R. W. K. and Lo, Y. M. D. (2017). Integrative single-cell and cell-free plasma RNA transcriptomics elucidates placental cellular dynamics. Proc Natl Acad Sci U S A 114, E7786–E7795.10.1073/pnas.1710470114 PMC5604038

50. van Beekhuizen, H. J., Joosten, I., de Groot, A. N., Lotgering, F. K., van der Laak, J. and Bulten, J. (2009). The number of multinucleated trophoblastic giant cells in the basal decidua is decreased in retained placenta. J Clin Pathol 62, 794–797.10.1136/jcp.2009.065953

51. VanOudenhove, J. J., Medina, R., Ghule, P. N., Lian, J. B., Stein, J. L., Zaidi, S. K. and Stein, G. S. (2016). Transient RUNX1 Expression during Early Mesendodermal Differentiation of hESCs Promotes Epithelial to Mesenchymal Transition through TGFB2 Signaling. Stem Cell Reports 7, 884–896.10.1016/j.stemcr.2016.09.006 PMC5106514

52. Vicovac, L. and Aplin, J. D. (1996). Epithelial-mesenchymal transition during trophoblast differentiation. Acta Anat (Basel) 156, 202–216

53. von Rango, U., Krusche, C. A., Kertschanska, S., Alfer, J., Kaufmann, P. and Beier, H. M. (2003). Apoptosis of extravillous trophoblast cells limits the trophoblast invasion in uterine but not in tubal pregnancy during first trimester. Placenta 24, 929–940

54. Wang, C., Wang, Y., Fu, Z., Huang, W., Yu, Z., Wang, J., Zheng, K., Zhang, S., Li, S. and Chen, J. (2022). MiR-29b-3p Inhibits Migration and Invasion of Papillary Thyroid Carcinoma by Downregulating COL1A1 and COL5A1. Front Oncol 12, 837581.10.3389/fonc.2022.837581 PMC9075584

55. Weiss, G., Sundl, M., Glasner, A., Huppertz, B. and Moser, G. (2016). The trophoblast plug during early pregnancy: a deeper insight. Histochem Cell Biol 146, 749–756.10.1007/s00418-016-1474-z PMC5101277

56. Yang, J., Antin, P., Berx, G., Blanpain, C., Brabletz, T., Bronner, M., Campbell, K., Cano, A., Casanova, J., Christofori, G., et al. (2020). Guidelines and definitions for research on epithelial-mesenchymal transition. Nat Rev Mol Cell Biol 21, 341–352.10.1038/s41580-020-0237-9 PMC7250738

57. Yang, X., Han, H., De Carvalho, D. D., Lay, F. D., Jones, P. A. and Liang, G. (2014). Gene body methylation can alter gene expression and is a therapeutic target in cancer. Cancer Cell 26, 577–590.10.1016/j.ccr.2014.07.028 PMC4224113

58. Yu, G., Wang, L. G., Han, Y. and He, Q. Y. (2012). clusterProfiler: an R package for comparing biological themes among gene clusters. OMICS 16, 284–287.10.1089/omi.2011.0118 PMC3339379

59. Zhang, B., Kim, M. Y., Elliot, G., Zhou, Y., Zhao, G., Li, D., Lowdon, R. F., Gormley, M., Kapidzic, M., Robinson, J. F., et al. (2021). Human placental cytotrophoblast epigenome dynamics over gestation and alterations in placental disease. Dev Cell 56, 1238–1252 e1235.10.1016/j.devcel.2021.04.001 PMC8650129

60. Zhao, M., Kong, L., Liu, Y. and Qu, H. (2015). dbEMT: an epithelial-mesenchymal transition associated gene resource. Sci Rep 5, 11459.10.1038/srep11459 PMC4477208

