## Supplementary tables S1-S5 for "Epigenetic changes regulating the epithelial-mesenchymal transition in human trophoblast differentiation": Supplementary_Tables_R1.docx

**Supplementary Table S1**:	Demographic characteristics of pregnancies from which placental tissue was obtained

| **Parameter** | **Full sample set (n=9)** | **EPIC set (n=4)** |
| --- | --- | --- |
| Maternal age (yrs) | 32.8 ± 1.2 | 33.5 ± 1.8 |
| Pre-pregnancy BMI | 25.8 ± 1.7 | 24.8 ± 2.6 |
| Gestational age at delivery (wks) | 38.9 ± 0.1 | 39 ± 01 |
| Fetal sex | M4 / F5 | M2 / F2 |
| Birthweight (g) | 3383 ± 92 | 3295 ± 58 |
| Birthweight centile | 56.8 ± 7.2 | 49.5 ± 4.1 |
| Placental weight (g) | 514 ± 17 | 483 ± 14 |

**Supplementary Table S2**:	EMT-associated genes showing differential expression between CTB and EVT

| **Gene ID** | **Fold-change** | **Gene ID** | **Fold-change** | **Gene ID** | **Fold-change** | **Gene ID** | **Fold-change** |
| --- | --- | --- | --- | --- | --- | --- | --- |
| MMP11 | 81.05 | RHOD | 8.07 | F11R | 2.98 | YAP1 | -2.70 |
| HLA-G | 78.18 | NOTCH2 | 7.88 | MAPK3 | 2.96 | ITGAV | -2.72 |
| PAPPA | 75.95 | MBP | 7.57 | CAPN2 | 2.95 | MME | -2.72 |
| CSH1 | 70.45 | RUNX1 | 7.54 | CD274 | 2.94 | ERF | -2.77 |
| RCN3 | 68.09 | FSTL1 | 7.45 | MYCN | 2.92 | TPD52L1 | -2.87 |
| PAPPA2 | 62.73 | TNF | 7.23 | STAT3 | 2.83 | TMEM30B | -2.96 |
| MCAM | 61.15 | TBX2 | 7.15 | GATA3 | 2.76 | CTNNBIP1 | -2.97 |
| SERPINE2 | 56.52 | FBLN5 | 6.99 | PTK2 | 2.76 | NUDT13 | -2.98 |
| B3GNT7 | 53.96 | CDR2 | 6.88 | CARD9 | 2.74 | NOTCH1 | -2.99 |
| FGFBP1 | 50.31 | PCOLCE | 6.88 | SEMA4C | 2.73 | MAP3K4 | -3.01 |
| VWCE | 45.29 | NDRG1 | 6.86 | YBX1 | 2.70 | DSC2 | -3.05 |
| PLAC8 | 42.85 | VCAN | 6.80 | MAP1B | 2.66 | TGM2 | -3.08 |
| PSG2 | 40.70 | COL6A3 | 6.77 | IL1B | 2.61 | RGS3 | -3.13 |
| NOG | 37.96 | SNAI1 | 6.72 | WNT5B | 2.58 | MTUS1 | -3.16 |
| ADAM19 | 37.63 | PTGS2 | 6.36 | TAGLN | 2.56 | PLAC4 | -3.21 |
| COL11A1 | 36.29 | FSCN1 | 6.22 | AXIN1 | 2.55 | STAT5B | -3.23 |
| SLPI | 35.76 | WWTR1 | 6.21 | FBN1 | 2.50 | RGS2 | -3.25 |
| FN1 | 35.59 | PRKCE | 6.01 | IFI30 | 2.45 | ADRB2 | -3.26 |
| LRRC15 | 32.61 | CTSZ | 5.82 | ID1 | 2.44 | CDK14 | -3.27 |
| ITGA5 | 29.39 | LUM | 5.76 | PLXND1 | 2.38 | GMNN | -3.28 |
| FGFR1 | 29.33 | KRT15 | 5.68 | SYT11 | 2.31 | IL18 | -3.28 |
| TGFB2 | 28.54 | DCN | 5.65 | FBLN1 | 2.31 | ECT2 | -3.39 |
| TIMP1 | 27.67 | CDH11 | 5.64 | FAP | 2.30 | VSNL1 | -3.40 |
| MTDH | 27.47 | CA2 | 5.58 | MMP14 | 2.28 | ZHX2 | -3.50 |
| LEP | 25.25 | HMGB3 | 5.55 | LRFN4 | 2.25 | SYK | -3.66 |
| LEP | 25.25 | LAD1 | 5.54 | POSTN | 2.23 | ERVH48-1 | -3.68 |
| SPARC | 24.26 | HPSE | 5.41 | GSN | 2.21 | KIT | -3.69 |
| MMP2 | 24.23 | CAV2 | 5.39 | ITGB3 | 2.21 | VPS13A | -3.73 |
| LHX2 | 23.78 | IGFBP4 | 5.36 | WT1 | 2.20 | CDKN2C | -3.77 |
| BGN | 23.77 | ERBB2 | 5.35 | HIF1A | 2.17 | RAPGEF5 | -3.85 |
| ADAM12 | 23.17 | GLRX | 5.33 | RAC1 | 2.16 | FAM169A | -3.86 |
| LIMS2 | 22.58 | L1CAM | 5.19 | RHOA | 2.13 | EGFR | -3.92 |
| TPM1 | 22.30 | PIK3CD | 5.19 | TNC | 2.11 | PTPRZ1 | -4.02 |
| IL1R1 | 22.26 | ZEB1 | 5.08 | MAPK8 | 2.03 | EML1 | -4.10 |
| NID2 | 20.54 | PDGFRB | 5.05 | EPHA1 | 1.95 | ITGA6 | -4.17 |
| RECK | 20.50 | TEAD1 | 4.94 | VEGFA | 1.90 | IGF1R | -4.20 |
| CSH2 | 18.80 | DAPK1 | 4.90 | CALD1 | 1.87 | SMAD7 | -4.22 |
| ENG | 18.69 | EMP3 | 4.89 | CXCL16 | 1.85 | GLT8D2 | -4.41 |
| MUC1 | 18.60 | DDR2 | 4.75 | TCF3 | 1.82 | CMTM8 | -4.48 |
| LOXL1 | 18.16 | TUBA1A | 4.63 | EIF5A2 | 1.82 | KRAS | -4.52 |
| COL6A1 | 18.05 | THY1 | 4.50 | MSN | 1.81 | COL5A2 | -4.81 |
| COPZ2 | 17.97 | COL3A1 | 4.48 | PPARG | 1.80 | ITGB4 | -4.90 |
| SDC2 | 17.77 | EDNRB | 4.40 | LATS1 | 1.80 | LAMA1 | -4.90 |
| HPGD | 17.59 | PTHLH | 4.38 | GAB2 | 1.77 | SULF1 | -4.98 |
| TGFB1 | 17.46 | TXNIP | 4.38 | WNT11 | 1.76 | PLS1 | -5.43 |
| TGFB1 | 17.46 | EPAS1 | 4.38 | PKP3 | 1.74 | EZH2 | -5.46 |
| SMAD3 | 17.23 | CXCR4 | 4.32 | S100P | 1.70 | RAB31 | -5.49 |
| SLC39A6 | 16.78 | SPOCK1 | 4.29 | CTSK | 1.70 | PLAU | -5.88 |
| TIMP3 | 16.01 | GALNT10 | 4.26 | SDC1 | 1.70 | VDR | -6.71 |
| PDE4A | 15.76 | SRC | 4.26 | CSK | 1.67 | TWIST1 | -6.80 |
| PSG5 | 15.49 | LIMA1 | 4.23 | AXL | 1.67 | BCL2 | -6.85 |
| SERPINE1 | 15.38 | SOD2 | 4.23 | TGFBR3 | 1.65 | CDS1 | -7.25 |
| SLC16A3 | 14.88 | IL6R | 4.19 | MDK | 1.64 | ERVW-1 | -7.41 |
| COL6A2 | 14.27 | MYL9 | 4.19 | JUP | 1.61 | MET | -7.75 |
| FLT4 | 13.93 | FHL2 | 4.15 | EGR1 | 1.58 | SEMA3E | -8.26 |
| CAMK2N1 | 13.90 | PRSS8 | 4.03 | HDGF | 1.57 | ERVV-2 | -8.47 |
| S100A4 | 13.90 | HOXB7 | 4.03 | HMOX1 | 1.57 | EPCAM | -8.62 |
| LGALS1 | 13.66 | VIM | 4.01 | ANPEP | 1.56 | PLXNB1 | -8.62 |
| RUSC2 | 13.61 | PSG4 | 3.88 | STAT5A | 1.55 | HBEGF | -8.85 |
| IGFBP3 | 13.55 | SPP1 | 3.84 | HS3ST3B1 | 1.54 | AURKA | -10.64 |
| FLT1 | 13.33 | BMP1 | 3.80 | MYC | -1.51 | CDKN2A | -10.99 |
| RGS4 | 13.31 | PRKCA | 3.79 | TP53 | -1.52 | PARP1 | -11.49 |
| NRN1 | 13.09 | KLF8 | 3.74 | DDX5 | -1.53 | FOXC1 | -12.05 |
| LOXL2 | 12.73 | FGFR3 | 3.71 | JAG2 | -1.56 | ID2 | -12.20 |
| HSD3B1 | 12.70 | ITGB1 | 3.70 | BCL2L1 | -1.58 | TBX3 | -12.99 |
| LTBP1 | 12.64 | TIAM1 | 3.70 | ITGBL1 | -1.59 | FXYD3 | -13.16 |
| MICALL2 | 12.55 | PRRG4 | 3.63 | MAP7 | -1.61 | FOXM1 | -13.89 |
| PAG1 | 12.54 | KRT17 | 3.58 | PIM3 | -1.62 | SPINT1 | -14.29 |
| PSG11 | 11.92 | MID1 | 3.54 | TCF7 | -1.68 | ALDH1A3 | -14.71 |
| TPM2 | 11.19 | CYP1B1 | 3.54 | SLC7A5 | -1.68 | DAB2 | -15.15 |
| CRISPLD2 | 11.19 | ESR1 | 3.54 | ELF3 | -1.69 | JAG1 | -15.87 |
| GJB3 | 11.05 | COL1A2 | 3.53 | HMGB1 | -1.72 | ERVV-1 | -16.39 |
| FST | 10.64 | CLU | 3.51 | NF1 | -1.80 | CD24 | -16.67 |
| NUAK1 | 10.19 | AEBP1 | 3.45 | CBR1 | -1.86 | CLDN1 | -18.87 |
| LPCAT1 | 10.18 | TBX20 | 3.44 | MUC16 | -1.88 | FGFR2 | -19.23 |
| TFPI | 10.06 | LTBP2 | 3.42 | KLF13 | -1.95 | ANK3 | -21.74 |
| MFAP5 | 10.06 | TCF4 | 3.39 | FZD7 | -1.98 | BMP7 | -23.81 |
| SMAD9 | 10.06 | OVOL2 | 3.38 | SCRIB | -2.06 | MARVELD3 | -28.57 |
| PSG3 | 10.00 | PLAT | 3.37 | CDKN1B | -2.11 | OCLN | -43.48 |
| SYNE1 | 9.95 | KRT14 | 3.36 | COL8A1 | -2.15 | MSX2 | -47.62 |
| KIF3C | 9.94 | KLF6 | 3.34 | IDH1 | -2.23 | SERPINF1 | -52.63 |
| COL5A1 | 9.86 | KLF6 | 3.34 | BRAF | -2.24 | ST14 | -71.43 |
| PLAUR | 9.73 | HSPB1 | 3.32 | LAMA5 | -2.24 | NRP1 | -83.33 |
| PTN | 9.35 | JAK2 | 3.26 | LYPD3 | -2.28 | PROM1 | -83.33 |
| PTN | 9.35 | KRT7 | 3.21 | PPL | -2.38 | CYP19A1 | -100.00 |
| ETV4 | 9.31 | ST6GALNAC2 | 3.18 | PTPN14 | -2.43 | SIGLEC6 | -111.11 |
| EPYC | 8.85 | LIMS1 | 3.15 | AGER | -2.44 | ELF5 | -125.00 |
| PHLDA1 | 8.83 | CAV1 | 3.08 | MAPK14 | -2.52 | NRP2 | -125.00 |
| PRRX1 | 8.34 | ABR | 3.08 | ZNF165 | -2.53 | ERVFRD-1 | -142.86 |
| PMP22 | 8.31 | DLX4 | 3.06 | HRAS | -2.60 | HS6ST2 | -142.86 |
| KRT19 | 8.28 | DLX4 | 3.06 | THBD | -2.64 | INSL4 | -142.86 |
| NFATC1 | 8.13 | PKP2 | 3.03 | CTNNB1 | -2.69 | PEG10 | -200.00 |
| FOXQ1 | 8.12 | TFPI2 | 3.00 | EDN1 | -2.70 | SLC27A2 | -250.00 |

**Supplementary Table S3**:	EMT-associated genes showing differential expression between vCTB and CTB

| Gene ID | Fold Change |
| --- | --- |
| LUM | 8.22 |
| DCN | 7.68 |
| CYP1B1 | 7.56 |
| CDH11 | 7.30 |
| VIM | 5.75 |
| COL6A3 | 5.73 |
| ZEB1 | 5.45 |
| VCAN | 5.32 |
| COL6A2 | 3.93 |
| TIMP1 | 3.75 |
| COL3A1 | 2.98 |
| LGALS1 | 2.70 |
| SDC2 | 2.58 |
| IGFBP4 | 2.26 |
| TXNIP | 2.08 |
| S100A4 | 2.01 |
| ZYX | -1.88 |
| GSK3A | -1.98 |
| TCF7 | -2.00 |
| LAMA5 | -2.05 |
| GATA3 | -2.46 |
| DLX4 | -2.52 |
| EGR1 | -2.86 |
| NFIC | -3.30 |
| MUC16 | -3.69 |

**Supplementary Table S4**:	Differential expression between CTB and EVT of primarily placental EMT-associated genes

| Gene ID | Fold Change |
| --- | --- |
| CSH2 | 38.00 |
| CSH1 | 37.41 |
| CSHL1 | 25.91 |
| PAPPA | 22.96 |
| HLA-G | 21.39 |
| PAPPA2 | 18.63 |
| PLAC8 | 15.98 |
| GH2 | 9.65 |
| LEP | 9.00 |
| PSG11 | 7.77 |
| PSG5 | 6.42 |
| HSD3B1 | 5.67 |
| PSG8 | 5.41 |
| PTN | 4.46 |
| PSG3 | 3.91 |
| PSG7 | 2.63 |
| PSG1 | 2.40 |
| PSG9 | 2.29 |
| PLAC1 | 2.24 |
| PSG4 | 2.22 |
| EDNRB | 2.14 |
| PLAC9 | 2.02 |
| MID1 | 1.74 |
| PLAC2 | -1.87 |
| PLAC4 | -6.15 |
| ERVW-1 | -14.24 |
| ERVV-2 | -19.41 |
| ERVV-1 | -22.80 |
| CYP19A1 | -164.72 |
| SIGLEC6 | -213.61 |
| PEG10 | -329.45 |
| INSL4 | -332.75 |
| ERVFRD-1 | -343.84 |

**Supplementary Table S5**:	Potential effects on the EMT of altered expression of DE-EMT genes that show a gain of methylation

| **Gene ID** | **Fold change** | **Effect on EMT** | **References** |
| --- | --- | --- | --- |
| MUC1 | 18.60 | Pro-EMT | (Qing et al., 2022; Rajabi and Kufe, 2017) |
| TGFB1 | 17.46 | Pro-EMT | (Hao et al., 2019; Pang et al., 2016) |
| FLT4 | 13.93 | Pro-EMT | (Kong et al., 2021; Kurmyshkina et al., 2020) |
| LOXL2 | 12.73 | Pro-EMT | (Cuevas et al., 2017; Park et al., 2017) |
| MICALL2 | 12.54 | Pro-EMT | (Chen et al., 2023; Zhu et al., 2015)2 |
| LPCAT1 | 10.17 | Pro-EMT | (Bi et al., 2019; Shen et al., 2022) |
| COL5A1 | 9.85 | Pro-EMT | (Liu et al., 2018d; Tsai et al., 2021) |
| NFATC1 | 8.13 | Pro-EMT | (Shen et al., 2021; Singh et al., 2015) |
| RUNX1 | 7.54 | Pro-EMT/Pro-MET | (Khawaled and Aqeilan, 2017; Li et al., 2019b; Zhou et al., 2018) |
| FSCN1 | 6.22 | Pro-EMT | (Li et al., 2018; Li et al., 2022b) |
| TEAD1 | 4.94 | Pro-EMT/Pro-MET | (Cruz et al., 2024; Huh et al., 2019) |
| PRSS8 | 4.03 | Pro-MET | (Bao et al., 2019; Li et al., 2023) |
| BMP1 | 3.80 | Pro-EMT/Pro-MET | (Wan et al., 2024; Zhu et al., 2020) |
| FGFR3 | 3.71 | Pro-EMT | (Li et al., 2019a; Zheng et al., 2023) |
| TIAM1 | 3.70 | Pro-EMT | (Ginn et al., 2023; Liu et al., 2018c) |
| AEBP1 | 3.45 | Pro-EMT | (Li et al., 2021; Liu et al., 2018b) |
| ABR | 3.08 | Pro-EMT | (Ungefroren et al., 2018; Vaughan et al., 2011) |
| PKP2 | 3.03 | Pro-EMT | (Rickelt, 2012; Wu et al., 2021) |
| PLXND1 | 2.38 | Pro-EMT | (Foley et al., 2015; Hagihara et al., 2022) |
| PKP3 | 1.74 | Pro-EMT/Pro-MET | (Aigner et al., 2007; Liu et al., 2024) |
| CSK | 1.67 | Pro-EMT | (Li et al., 2019c; Ortiz et al., 2021) |
| TGFBR3 | 1.65 | Pro-EMT/Pro-MET | (Lu et al., 2023; Nishida et al., 2018) |
| BCL2L1 | -1.58 | Pro-MET | (Keitel et al., 2014; Zhang et al., 2023) |
| SLC7A5 | -1.68 | Pro-MET | (Liu et al., 2022; Tornroos et al., 2022) |
| KLF13 | -1.95 | Pro-MET | (Chen et al., 2022; Li et al., 2022a; Liu et al., 2023) |
| PTPN14 | -2.43 | Pro-EMT | (Belle et al., 2015; Liu et al., 2013) |
| HRAS | -2.60 | Pro-MET | (He et al., 2015a; Wong et al., 2013) |
| MME | -2.72 | Pro-MET | (Imokawa, 2016; Song et al., 2016) |
| TPD52L1 | -2.88 | Pro-MET | (Boutros and Byrne, 2005; Hong et al., 2021) |
| CTNNBIP1 | -2.97 | Pro-EMT | (Bi et al., 2018; Chang et al., 2019) |
| NOTCH1 | -2.99 | Pro-MET | (Natsuizaka et al., 2017; Shao et al., 2015)3 |
| MAP3K4 | -3.01 | Pro-EMT | (Abell et al., 2011; Raghu et al., 2019) |
| TGM2 | -3.08 | Pro-MET | (He et al., 2015b; Ma et al., 2018) |
| CDK14 | -3.27 | Pro-MET | (Ou-Yang et al., 2017; Zhang et al., 2022) |
| VSNL1 | 0.2939 | Pro-MET | (Aiba et al., 2023; Dai et al., 2020) |
| SYK | 0.2727 | Pro-MET/Pro-EMT | (Geahlen, 2014; Krisenko and Geahlen, 2015) |
| EGFR | 0.2554 | Pro-MET/Pro-EMT | (Lo et al., 2007; Schinke et al., 2022) |
| EML1 | 0.2442 | Pro-MET | (Markus et al., 2021; Poria et al., 2022) |
| IGF1R | 0.2383 | Pro-MET | (Alfaro-Arnedo et al., 2022; Vazquez-Martin et al., 2013) |
| COL5A2 | 0.2077 | Pro-MET | (Jin et al., 2023; Wu et al., 2022) |
| ITGB4 | 0.2040 | Pro-MET | (Li et al., 2017; Masugi et al., 2015) |
| BCL2 | 0.1458 | Pro-MET | (Du et al., 2018; Zuo et al., 2010) |
| MET | 0.1294 | Pro-MET | (Wang et al., 2021; Zhang et al., 2018) |
| PLXNB1 | 0.1162 | Pro-MET | (Casazza et al., 2010; Tam et al., 2017) |
| FXYD3 | 0.0761 | Pro-MET/Pro-EMT | (Kayed et al., 2006; Yamamoto et al., 2011) |
| SPINT1 | 0.0698 | Pro-MET/Pro-EMT | (Cheng et al., 2009; Gomez-Abenza et al., 2019; Liu et al., 2018a) |
| ALDH1A3 | 0.0676 | Pro-MET | (McLean et al., 2023; Yamashita et al., 2020) |
| JAG1 | 0.0635 | Pro-MET | (Negri et al., 2023; Xiu et al., 2020) |
| FGFR2 | 0.0523 | Pro-MET/Pro-EMT | (Grygielewicz et al., 2016; Lei et al., 2021; Xu et al., 2022b) |
| ANK3 | 0.0459 | Pro-EMT | (Somsuan and Aluksanasuwan, 2023; Zeng et al., 2022) |
| BMP7 | 0.0416 | Pro-MET/Pro-EMT | (Sun et al., 2020; Ying et al., 2015; Zeisberg et al., 2003) |
| MSX2 | 0.0214 | Pro-MET | (Liang et al., 2016; Zhai et al., 2011) |
| ELF5 | 0.0084 | Pro-EMT | (Chakrabarti et al., 2012; Wu et al., 2015) |
| NRP2 | 0.0081 | Pro-MET | (Gemmill et al., 2017; Islam et al., 2022) |
| SLC27A2 | 0.0040 | Pro-MET/Pro-EMT | (Veglia et al., 2019; Xu et al., 2022a) |

Pro-EMT: Change which has the effect of promoting the EMT/suppressing MET

Pro-MET: Change which has the effect of suppressing the EMT/promoting MET

Pro-EMT/Pro-MET: May have the effect of promoting EMT or MET, depending on context

References

**Abell, A. N., Jordan, N. V., Huang, W., Prat, A., Midland, A. A., Johnson, N. L., Granger, D. A., Mieczkowski, P. A., Perou, C. M., Gomez, S. M., et al.** (2011). MAP3K4/CBP-regulated H2B acetylation controls epithelial-mesenchymal transition in trophoblast stem cells. *Cell Stem Cell* **8**, 525-537.

**Aiba, T., Hijiya, N., Akagi, T., Tsukamoto, Y., Hirashita, Y., Kinoshita, K., Uchida, T., Nakada, C., Kurogi, S., Ueda, Y., et al.** (2023). Overexpression of VSNL1 enhances cell proliferation in colorectal carcinogenesis. *Pathobiology*.

**Aigner, K., Descovich, L., Mikula, M., Sultan, A., Dampier, B., Bonne, S., van Roy, F., Mikulits, W., Schreiber, M., Brabletz, T., et al.** (2007). The transcription factor ZEB1 (deltaEF1) represses Plakophilin 3 during human cancer progression. *FEBS Lett* **581**, 1617-1624.

**Alfaro-Arnedo, E., Lopez, I. P., Pineiro-Hermida, S., Canalejo, M., Gotera, C., Sola, J. J., Roncero, A., Peces-Barba, G., Ruiz-Martinez, C. and Pichel, J. G.** (2022). IGF1R acts as a cancer-promoting factor in the tumor microenvironment facilitating lung metastasis implantation and progression. *Oncogene* **41**, 3625-3639.

**Bao, Y., Guo, Y., Yang, Y., Wei, X., Zhang, S., Zhang, Y., Li, K., Yuan, M., Guo, D., Macias, V., et al.** (2019). PRSS8 suppresses colorectal carcinogenesis and metastasis. *Oncogene* **38**, 497-517.

**Belle, L., Ali, N., Lonic, A., Li, X., Paltridge, J. L., Roslan, S., Herrmann, D., Conway, J. R., Gehling, F. K., Bert, A. G., et al.** (2015). The tyrosine phosphatase PTPN14 (Pez) inhibits metastasis by altering protein trafficking. *Sci Signal* **8**, ra18.

**Bi, J., Ichu, T. A., Zanca, C., Yang, H., Zhang, W., Gu, Y., Chowdhry, S., Reed, A., Ikegami, S., Turner, K. M., et al.** (2019). Oncogene Amplification in Growth Factor Signaling Pathways Renders Cancers Dependent on Membrane Lipid Remodeling. *Cell Metab* **30**, 525-538 e528.

**Bi, W., Huang, J., Nie, C., Liu, B., He, G., Han, J., Pang, R., Ding, Z., Xu, J. and Zhang, J.** (2018). CircRNA circRNA_102171 promotes papillary thyroid cancer progression through modulating CTNNBIP1-dependent activation of beta-catenin pathway. *J Exp Clin Cancer Res* **37**, 275.

**Boutros, R. and Byrne, J. A.** (2005). D53 (TPD52L1) is a cell cycle-regulated protein maximally expressed at the G2-M transition in breast cancer cells. *Exp Cell Res* **310**, 152-165.

**Casazza, A., Finisguerra, V., Capparuccia, L., Camperi, A., Swiercz, J. M., Rizzolio, S., Rolny, C., Christensen, C., Bertotti, A., Sarotto, I., et al.** (2010). Sema3E-Plexin D1 signaling drives human cancer cell invasiveness and metastatic spreading in mice. *J Clin Invest* **120**, 2684-2698.

**Chakrabarti, R., Hwang, J., Andres Blanco, M., Wei, Y., Lukacisin, M., Romano, R. A., Smalley, K., Liu, S., Yang, Q., Ibrahim, T., et al.** (2012). Elf5 inhibits the epithelial-mesenchymal transition in mammary gland development and breast cancer metastasis by transcriptionally repressing Snail2. *Nat Cell Biol* **14**, 1212-1222.

**Chang, J. M., Tsai, A. C., Huang, W. R. and Tseng, R. C.** (2019). The Alteration of CTNNBIP1 in Lung Cancer. *Int J Mol Sci* **20**.

**Chen, C. C., Xie, X. M., Zhao, X. K., Zuo, S. and Li, H. Y.** (2022). Kruppel-like Factor 13 Promotes HCC Progression by Transcriptional Regulation of HMGCS1-mediated Cholesterol Synthesis. *J Clin Transl Hepatol* **10**, 1125-1137.

**Chen, Y., Song, W., Zhang, H. and Ji, X.** (2023). MICALL2 participates in the regulation of epithelial-mesenchymal transition in alveolar epithelial cells - Potential roles in pulmonary fibrosis. *Arch Biochem Biophys* **747**, 109730.

**Cheng, H., Fukushima, T., Takahashi, N., Tanaka, H. and Kataoka, H.** (2009). Hepatocyte growth factor activator inhibitor type 1 regulates epithelial to mesenchymal transition through membrane-bound serine proteinases. *Cancer Res* **69**, 1828-1835.

**Cruz, S. P., Zhang, Q., Devarajan, R., Paia, C., Luo, B., Zhang, K., Koivusalo, S., Qin, L., Xia, J., Ahtikoski, A., et al.** (2024). Dampened Regulatory Circuitry of TEAD1/ITGA1/ITGA2 Promotes TGFbeta1 Signaling to Orchestrate Prostate Cancer Progression. *Adv Sci (Weinh)*, e2305547.

**Cuevas, E. P., Eraso, P., Mazon, M. J., Santos, V., Moreno-Bueno, G., Cano, A. and Portillo, F.** (2017). LOXL2 drives epithelial-mesenchymal transition via activation of IRE1-XBP1 signalling pathway. *Sci Rep* **7**, 44988.

**Dai, Q. Q., Wang, Y. Y., Jiang, Y. P., Li, L. and Wang, H. J.** (2020). VSNL1 Promotes Gastric Cancer Cell Proliferation and Migration by Regulating P2X3/P2Y2 Receptors and Is a Clinical Indicator of Poor Prognosis in Gastric Cancer Patients. *Gastroenterol Res Pract* **2020**, 7241942.

**Du, C., Zhang, X., Yao, M., Lv, K., Wang, J., Chen, L., Chen, Y., Wang, S. and Fu, P.** (2018). Bcl-2 promotes metastasis through the epithelial-to-mesenchymal transition in the BCap37 medullary breast cancer cell line. *Oncol Lett* **15**, 8991-8898.

**Foley, K., Rucki, A. A., Xiao, Q., Zhou, D., Leubner, A., Mo, G., Kleponis, J., Wu, A. A., Sharma, R., Jiang, Q., et al.** (2015). Semaphorin 3D autocrine signaling mediates the metastatic role of annexin A2 in pancreatic cancer. *Sci Signal* **8**, ra77.

**Geahlen, R. L.** (2014). Getting Syk: spleen tyrosine kinase as a therapeutic target. *Trends Pharmacol Sci* **35**, 414-422.

**Gemmill, R. M., Nasarre, P., Nair-Menon, J., Cappuzzo, F., Landi, L., D'Incecco, A., Uramoto, H., Yoshida, T., Haura, E. B., Armeson, K. and Drabkin, H. A.** (2017). The neuropilin 2 isoform NRP2b uniquely supports TGFbeta-mediated progression in lung cancer. *Sci Signal* **10**.

**Ginn, L., Maltas, J., Baker, M. J., Chaturvedi, A., Wilson, L., Guilbert, R., Amaral, F. M. R., Priest, L., Mole, H., Blackhall, F., et al.** (2023). A TIAM1-TRIM28 complex mediates epigenetic silencing of protocadherins to promote migration of lung cancer cells. *Proc Natl Acad Sci U S A* **120**, e2300489120.

**Gomez-Abenza, E., Ibanez-Molero, S., Garcia-Moreno, D., Fuentes, I., Zon, L. I., Mione, M. C., Cayuela, M. L., Gabellini, C. and Mulero, V.** (2019). Zebrafish modeling reveals that SPINT1 regulates the aggressiveness of skin cutaneous melanoma and its crosstalk with tumor immune microenvironment. *J Exp Clin Cancer Res* **38**, 405.

**Grygielewicz, P., Dymek, B., Bujak, A., Gunerka, P., Stanczak, A., Lamparska-Przybysz, M., Wieczorek, M., Dzwonek, K. and Zdzalik, D.** (2016). Epithelial-mesenchymal transition confers resistance to selective FGFR inhibitors in SNU-16 gastric cancer cells. *Gastric Cancer* **19**, 53-62.

**Hagihara, K., Haraguchi, N., Nishimura, J., Yasueda, A., Fujino, S., Ogino, T., Takahashi, H., Miyoshi, N., Uemura, M., Matsuda, C., et al.** (2022). PLXND1/SEMA3E Promotes Epithelial-Mesenchymal Transition Partly via the PI3K/AKT-Signaling Pathway and Induces Heterogenity in Colorectal Cancer. *Ann Surg Oncol* **29**, 7435-7445.

**Hao, Y., Baker, D. and Ten Dijke, P.** (2019). TGF-beta-Mediated Epithelial-Mesenchymal Transition and Cancer Metastasis. *Int J Mol Sci* **20**.

**He, F., Melamed, J., Tang, M. S., Huang, C. and Wu, X. R.** (2015a). Oncogenic HRAS Activates Epithelial-to-Mesenchymal Transition and Confers Stemness to p53-Deficient Urothelial Cells to Drive Muscle Invasion of Basal Subtype Carcinomas. *Cancer Res* **75**, 2017-2028.

**He, W., Sun, Z. and Liu, Z.** (2015b). Silencing of TGM2 reverses epithelial to mesenchymal transition and modulates the chemosensitivity of breast cancer to docetaxel. *Exp Ther Med* **10**, 1413-1418.

**Hong, Q., Li, B., Cai, X., Lv, Z., Cai, S., Zhong, Y. and Wen, B.** (2021). Transcriptomic Analyses of the Adenoma-Carcinoma Sequence Identify Hallmarks Associated With the Onset of Colorectal Cancer. *Front Oncol* **11**, 704531.

**Huh, H. D., Kim, D. H., Jeong, H. S. and Park, H. W.** (2019). Regulation of TEAD Transcription Factors in Cancer Biology. *Cells* **8**.

**Imokawa, G.** (2016). Epithelial-mesenchymal interaction mechanisms leading to the over-expression of neprilysin are involved in the UVB-induced formation of wrinkles in the skin. *Exp Dermatol* **25 Suppl 3**, 2-13.

**Islam, R., Mishra, J., Bodas, S., Bhattacharya, S., Batra, S. K., Dutta, S. and Datta, K.** (2022). Role of Neuropilin-2-mediated signaling axis in cancer progression and therapy resistance. *Cancer Metastasis Rev* **41**, 771-787.

**Jin, Y., Song, X., Sun, X. and Ding, Y.** (2023). Up-regulation of collagen type V alpha 2 (COL5A2) promotes malignant phenotypes in gastric cancer cell via inducing epithelial-mesenchymal transition (EMT). *Open Med (Wars)* **18**, 20220593.

**Kayed, H., Kleeff, J., Kolb, A., Ketterer, K., Keleg, S., Felix, K., Giese, T., Penzel, R., Zentgraf, H., Buchler, M. W., et al.** (2006). FXYD3 is overexpressed in pancreatic ductal adenocarcinoma and influences pancreatic cancer cell growth. *Int J Cancer* **118**, 43-54.

**Keitel, U., Scheel, A., Thomale, J., Halpape, R., Kaulfuss, S., Scheel, C. and Dobbelstein, M.** (2014). Bcl-xL mediates therapeutic resistance of a mesenchymal breast cancer cell subpopulation. *Oncotarget* **5**, 11778-11791.

**Khawaled, S. and Aqeilan, R. I.** (2017). RUNX1, a new regulator of EMT in breast cancer. *Oncotarget* **8**, 17407-17408.

**Kong, D., Zhou, H., Neelakantan, D., Hughes, C. J., Hsu, J. Y., Srinivasan, R. R., Lewis, M. T. and Ford, H. L.** (2021). VEGF-C mediates tumor growth and metastasis through promoting EMT-epithelial breast cancer cell crosstalk. *Oncogene* **40**, 964-979.

**Krisenko, M. O. and Geahlen, R. L.** (2015). Calling in SYK: SYK's dual role as a tumor promoter and tumor suppressor in cancer. *Biochim Biophys Acta* **1853**, 254-263.

**Kurmyshkina, O., Kovchur, P., Schegoleva, L. and Volkova, T.** (2020). Markers of Angiogenesis, Lymphangiogenesis, and Epithelial-Mesenchymal Transition (Plasticity) in CIN and Early Invasive Carcinoma of the Cervix: Exploring Putative Molecular Mechanisms Involved in Early Tumor Invasion. *Int J Mol Sci* **21**.

**Lei, J. H., Lee, M. H., Miao, K., Huang, Z., Yao, Z., Zhang, A., Xu, J., Zhao, M., Huang, Z., Zhang, X., et al.** (2021). Activation of FGFR2 Signaling Suppresses BRCA1 and Drives Triple-Negative Mammary Tumorigenesis That is Sensitive to Immunotherapy. *Adv Sci (Weinh)* **8**, e2100974.

**Li, B., Pang, S., Dou, J., Zhou, C., Shen, B. and Zhou, Y.** (2022a). The inhibitory effect of LINC00261 upregulation on the pancreatic cancer EMT process is mediated by KLF13 via the mTOR signaling pathway. *Clin Transl Oncol* **24**, 1059-1072.

**Li, D., Liu, Z., Ding, X. and Qin, Z.** (2021). AEBP1 Is One of the Epithelial-Mesenchymal Transition Regulatory Genes in Colon Adenocarcinoma. *Biomed Res Int* **2021**, 3108933.

**Li, D., Xia, L., Huang, P., Wang, Z., Guo, Q., Huang, C., Leng, W. and Qin, S.** (2023). Heterogeneity and plasticity of epithelial-mesenchymal transition (EMT) in cancer metastasis: Focusing on partial EMT and regulatory mechanisms. *Cell Prolif* **56**, e13423.

**Li, J., Zhang, S., Pei, M., Wu, L., Liu, Y., Li, H., Lu, J. and Li, X.** (2018). FSCN1 Promotes Epithelial-Mesenchymal Transition Through Increasing Snail1 in Ovarian Cancer Cells. *Cell Physiol Biochem* **49**, 1766-1777.

**Li, L., Zhang, S., Li, H. and Chou, H.** (2019a). FGFR3 promotes the growth and malignancy of melanoma by influencing EMT and the phosphorylation of ERK, AKT, and EGFR. *BMC Cancer* **19**, 963.

**Li, Q., Lai, Q., He, C., Fang, Y., Yan, Q., Zhang, Y., Wang, X., Gu, C., Wang, Y., Ye, L., et al.** (2019b). RUNX1 promotes tumour metastasis by activating the Wnt/beta-catenin signalling pathway and EMT in colorectal cancer. *J Exp Clin Cancer Res* **38**, 334.

**Li, X., Wang, F., Ren, M., Du, M. and Zhou, J.** (2019c). The effects of c-Src kinase on EMT signaling pathway in human lens epithelial cells associated with lens diseases. *BMC Ophthalmol* **19**, 219.

**Li, X. L., Liu, L., Li, D. D., He, Y. P., Guo, L. H., Sun, L. P., Liu, L. N., Xu, H. X. and Zhang, X. P.** (2017). Integrin beta4 promotes cell invasion and epithelial-mesenchymal transition through the modulation of Slug expression in hepatocellular carcinoma. *Sci Rep* **7**, 40464.

**Li, Z., Shi, J., Zhang, N., Zheng, X., Jin, Y., Wen, S., Hu, W., Wu, Y. and Gao, W.** (2022b). FSCN1 acts as a promising therapeutic target in the blockade of tumor cell motility: a review of its function, mechanism, and clinical significance. *J Cancer* **13**, 2528-2539.

**Liang, H., Zhang, Q., Lu, J., Yang, G., Tian, N., Wang, X., Tan, Y. and Tan, D.** (2016). MSX2 Induces Trophoblast Invasion in Human Placenta. *PLoS One* **11**, e0153656.

**Liu, B., Feng, Y., Xie, N., Yang, Y. and Yang, D.** (2024). FERMT1 promotes cell migration and invasion in non-small cell lung cancer via regulating PKP3-mediated activation of p38 MAPK signaling. *BMC Cancer* **24**, 58.

**Liu, C. L., Yang, P. S., Chien, M. N., Chang, Y. C., Lin, C. H. and Cheng, S. P.** (2018a). Expression of serine peptidase inhibitor Kunitz type 1 in differentiated thyroid cancer. *Histochem Cell Biol* **149**, 635-644.

**Liu, J. Y., Jiang, L., Liu, J. J., He, T., Cui, Y. H., Qian, F. and Yu, P. W.** (2018b). AEBP1 promotes epithelial-mesenchymal transition of gastric cancer cells by activating the NF-kappaB pathway and predicts poor outcome of the patients. *Sci Rep* **8**, 11955.

**Liu, L., Wu, B., Cai, H., Li, D., Ma, Y., Zhu, X., Lv, Z., Fan, Y. and Zhang, X.** (2018c). Tiam1 promotes thyroid carcinoma metastasis by modulating EMT via Wnt/beta-catenin signaling. *Exp Cell Res* **362**, 532-540.

**Liu, W., Wei, H., Gao, Z., Chen, G., Liu, Y., Gao, X., Bai, G., He, S., Liu, T., Xu, W., et al.** (2018d). COL5A1 may contribute the metastasis of lung adenocarcinoma. *Gene* **665**, 57-66.

**Liu, X., Yang, N., Figel, S. A., Wilson, K. E., Morrison, C. D., Gelman, I. H. and Zhang, J.** (2013). PTPN14 interacts with and negatively regulates the oncogenic function of YAP. *Oncogene* **32**, 1266-1273.

**Liu, Y., Ma, G., Liu, J., Zheng, H., Huang, G., Song, Q., Pang, Z. and Du, J.** (2022). SLC7A5 is a lung adenocarcinoma-specific prognostic biomarker and participates in forming immunosuppressive tumor microenvironment. *Heliyon* **8**, e10866.

**Liu, Y., Song, Y., He, Y., Kong, Z., Li, H., Zhu, Y. and Liu, S.** (2023). Kruppel-like factor 13 acts as a tumor suppressor in thyroid carcinoma by downregulating IFIT1. *Biol Direct* **18**, 65.

**Lo, H. W., Hsu, S. C., Xia, W., Cao, X., Shih, J. Y., Wei, Y., Abbruzzese, J. L., Hortobagyi, G. N. and Hung, M. C.** (2007). Epidermal growth factor receptor cooperates with signal transducer and activator of transcription 3 to induce epithelial-mesenchymal transition in cancer cells via up-regulation of TWIST gene expression. *Cancer Res* **67**, 9066-9076.

**Lu, P., Wu, B., Wang, Y., Russell, M., Liu, Y., Bernard, D. J., Zheng, D. and Zhou, B.** (2023). Prerequisite endocardial-mesenchymal transition for murine cardiac trabecular angiogenesis. *Dev Cell* **58**, 791-805 e794.

**Ma, H., Xie, L., Zhang, L., Yin, X., Jiang, H., Xie, X., Chen, R., Lu, H. and Ren, Z.** (2018). Activated hepatic stellate cells promote epithelial-to-mesenchymal transition in hepatocellular carcinoma through transglutaminase 2-induced pseudohypoxia. *Commun Biol* **1**, 168.

**Markus, F., Kannengiesser, A., Nader, P., Atigbire, P., Scholten, A., Vossing, C., Bultmann, E., Korenke, G. C., Owczarek-Lipska, M. and Neidhardt, J.** (2021). A novel missense variant in the EML1 gene associated with bilateral ribbon-like subcortical heterotopia leads to ciliary defects. *J Hum Genet* **66**, 1159-1167.

**Masugi, Y., Yamazaki, K., Emoto, K., Effendi, K., Tsujikawa, H., Kitago, M., Itano, O., Kitagawa, Y. and Sakamoto, M.** (2015). Upregulation of integrin beta4 promotes epithelial-mesenchymal transition and is a novel prognostic marker in pancreatic ductal adenocarcinoma. *Lab Invest* **95**, 308-319.

**McLean, M. E., MacLean, M. R., Cahill, H. F., Arun, R. P., Walker, O. L., Wasson, M. D., Fernando, W., Venkatesh, J. and Marcato, P.** (2023). The Expanding Role of Cancer Stem Cell Marker ALDH1A3 in Cancer and Beyond. *Cancers (Basel)* **15**.

**Natsuizaka, M., Whelan, K. A., Kagawa, S., Tanaka, K., Giroux, V., Chandramouleeswaran, P. M., Long, A., Sahu, V., Darling, D. S., Que, J., et al.** (2017). Interplay between Notch1 and Notch3 promotes EMT and tumor initiation in squamous cell carcinoma. *Nat Commun* **8**, 1758.

**Negri, F., Bottarelli, L., Pedrazzi, G., Maddalo, M., Leo, L., Milanese, G., Sala, R., Lecchini, M., Campanini, N., Bozzetti, C., et al.** (2023). Notch-Jagged1 signaling and response to bevacizumab therapy in advanced colorectal cancer: A glance to radiomics or back to physiopathology? *Front Oncol* **13**, 1132564.

**Nishida, J., Miyazono, K. and Ehata, S.** (2018). Decreased TGFBR3/betaglycan expression enhances the metastatic abilities of renal cell carcinoma cells through TGF-beta-dependent and -independent mechanisms. *Oncogene* **37**, 2197-2212.

**Ortiz, M. A., Mikhailova, T., Li, X., Porter, B. A., Bah, A. and Kotula, L.** (2021). Src family kinases, adaptor proteins and the actin cytoskeleton in epithelial-to-mesenchymal transition. *Cell Commun Signal* **19**, 67.

**Ou-Yang, J., Huang, L. H. and Sun, X. X.** (2017). Cyclin-Dependent Kinase 14 Promotes Cell Proliferation, Migration and Invasion in Ovarian Cancer by Inhibiting Wnt Signaling Pathway. *Gynecol Obstet Invest* **82**, 230-239.

**Pang, M. F., Georgoudaki, A. M., Lambut, L., Johansson, J., Tabor, V., Hagikura, K., Jin, Y., Jansson, M., Alexander, J. S., Nelson, C. M., et al.** (2016). TGF-beta1-induced EMT promotes targeted migration of breast cancer cells through the lymphatic system by the activation of CCR7/CCL21-mediated chemotaxis. *Oncogene* **35**, 748-760.

**Park, P. G., Jo, S. J., Kim, M. J., Kim, H. J., Lee, J. H., Park, C. K., Kim, H., Lee, K. Y., Kim, H., Park, J. H., et al.** (2017). Role of LOXL2 in the epithelial-mesenchymal transition and colorectal cancer metastasis. *Oncotarget* **8**, 80325-80335.

**Poria, D., Sun, C., Santeford, A., Kielar, M., Apte, R. S., Kisselev, O. G., Chen, S. and Kefalov, V. J.** (2022). EML1 is essential for retinal photoreceptor migration and survival. *Sci Rep* **12**, 2897.

**Qing, L., Li, Q. and Dong, Z.** (2022). MUC1: An emerging target in cancer treatment and diagnosis. *Bull Cancer* **109**, 1202-1216.

**Raghu, D., Mobley, R. J., Shendy, N. A. M., Perry, C. H. and Abell, A. N.** (2019). GALNT3 Maintains the Epithelial State in Trophoblast Stem Cells. *Cell Rep* **26**, 3684-3697 e3687.

**Rajabi, H. and Kufe, D.** (2017). MUC1-C Oncoprotein Integrates a Program of EMT, Epigenetic Reprogramming and Immune Evasion in Human Carcinomas. *Biochim Biophys Acta Rev Cancer* **1868**, 117-122.

**Rickelt, S.** (2012). Plakophilin-2: a cell-cell adhesion plaque molecule of selective and fundamental importance in cardiac functions and tumor cell growth. *Cell Tissue Res* **348**, 281-294.

**Schinke, H., Shi, E., Lin, Z., Quadt, T., Kranz, G., Zhou, J., Wang, H., Hess, J., Heuer, S., Belka, C., et al.** (2022). A transcriptomic map of EGFR-induced epithelial-to-mesenchymal transition identifies prognostic and therapeutic targets for head and neck cancer. *Mol Cancer* **21**, 178.

**Shao, S., Zhao, X., Zhang, X., Luo, M., Zuo, X., Huang, S., Wang, Y., Gu, S. and Zhao, X.** (2015). Notch1 signaling regulates the epithelial-mesenchymal transition and invasion of breast cancer in a Slug-dependent manner. *Mol Cancer* **14**, 28.

**Shen, L., Gu, P., Qiu, C., Ding, W. T., Zhang, L., Cao, W. Y., Li, Z. Y., Yan, B. and Sun, X.** (2022). Lysophosphatidylcholine acyltransferase 1 promotes epithelial-mesenchymal transition of hepatocellular carcinoma via the Wnt/beta-catenin signaling pathway. *Ann Hepatol* **27**, 100680.

**Shen, T., Yue, C., Wang, X., Wang, Z., Wu, Y., Zhao, C., Chang, P., Sun, X. and Wang, W.** (2021). NFATc1 promotes epithelial-mesenchymal transition and facilitates colorectal cancer metastasis by targeting SNAI1. *Exp Cell Res* **408**, 112854.

**Singh, S. K., Chen, N. M., Hessmann, E., Siveke, J., Lahmann, M., Singh, G., Voelker, N., Vogt, S., Esposito, I., Schmidt, A., et al.** (2015). Antithetical NFATc1-Sox2 and p53-miR200 signaling networks govern pancreatic cancer cell plasticity. *EMBO J* **34**, 517-530.

**Somsuan, K. and Aluksanasuwan, S.** (2023). Bioinformatic analyses reveal the prognostic significance and potential role of ankyrin 3 (ANK3) in kidney renal clear cell carcinoma. *Genomics Inform* **21**, e22.

**Song, S., Zhang, M., Yi, Z., Zhang, H., Shen, T., Yu, X., Zhang, C., Zheng, X., Yu, L., Ma, C., et al.** (2016). The role of PDGF-B/TGF-beta1/neprilysin network in regulating endothelial-to-mesenchymal transition in pulmonary artery remodeling. *Cell Signal* **28**, 1489-1501.

**Sun, R., Guan, H., Liu, W., Liang, J., Wang, F. and Li, C.** (2020). Expression of BMP7 in cervical cancer and inhibition of epithelial‑mesenchymal transition by BMP7 knockdown in HeLa cells. *Int J Mol Med* **45**, 1417-1424.

**Tam, K. J., Hui, D. H. F., Lee, W. W., Dong, M., Tombe, T., Jiao, I. Z. F., Khosravi, S., Takeuchi, A., Peacock, J. W., Ivanova, L., et al.** (2017). Semaphorin 3 C drives epithelial-to-mesenchymal transition, invasiveness, and stem-like characteristics in prostate cells. *Sci Rep* **7**, 11501.

**Tornroos, R., Tina, E. and Gothlin Eremo, A.** (2022). SLC7A5 is linked to increased expression of genes related to proliferation and hypoxia in estrogen‑receptor‑positive breast cancer. *Oncol Rep* **47**.

**Tsai, H. F., Chang, Y. C., Li, C. H., Chan, M. H., Chen, C. L., Tsai, W. C. and Hsiao, M.** (2021). Type V collagen alpha 1 chain promotes the malignancy of glioblastoma through PPRC1-ESM1 axis activation and extracellular matrix remodeling. *Cell Death Discov* **7**, 313.

**Ungefroren, H., Witte, D. and Lehnert, H.** (2018). The role of small GTPases of the Rho/Rac family in TGF-beta-induced EMT and cell motility in cancer. *Dev Dyn* **247**, 451-461.

**Vaughan, E. M., Miller, A. L., Yu, H. Y. and Bement, W. M.** (2011). Control of local Rho GTPase crosstalk by Abr. *Curr Biol* **21**, 270-277.

**Vazquez-Martin, A., Cufi, S., Oliveras-Ferraros, C., Torres-Garcia, V. Z., Corominas-Faja, B., Cuyas, E., Bonavia, R., Visa, J., Martin-Castillo, B., Barrajon-Catalan, E., et al.** (2013). IGF-1R/epithelial-to-mesenchymal transition (EMT) crosstalk suppresses the erlotinib-sensitizing effect of EGFR exon 19 deletion mutations. *Sci Rep* **3**, 2560.

**Veglia, F., Tyurin, V. A., Blasi, M., De Leo, A., Kossenkov, A. V., Donthireddy, L., To, T. K. J., Schug, Z., Basu, S., Wang, F., et al.** (2019). Fatty acid transport protein 2 reprograms neutrophils in cancer. *Nature* **569**, 73-78.

**Wan, S., Liu, X., Sun, R., Liu, H., Jiang, J. and Wu, B.** (2024). Activated hepatic stellate cell-derived Bmp-1 induces liver fibrosis via mediating hepatocyte epithelial-mesenchymal transition. *Cell Death Dis* **15**, 41.

**Wang, S., Ma, H., Yan, Y., Chen, Y., Fu, S., Wang, J., Wang, Y., Chen, H. and Liu, J.** (2021). cMET promotes metastasis and epithelial-mesenchymal transition in colorectal carcinoma by repressing RKIP. *J Cell Physiol* **236**, 3963-3978.

**Wong, C. E., Yu, J. S., Quigley, D. A., To, M. D., Jen, K. Y., Huang, P. Y., Del Rosario, R. and Balmain, A.** (2013). Inflammation and Hras signaling control epithelial-mesenchymal transition during skin tumor progression. *Genes Dev* **27**, 670-682.

**Wu, B., Cao, X., Liang, X., Zhang, X., Zhang, W., Sun, G. and Wang, D.** (2015). Epigenetic regulation of Elf5 is associated with epithelial-mesenchymal transition in urothelial cancer. *PLoS One* **10**, e0117510.

**Wu, L., Amjad, S., Yun, H., Mani, S. and de Perrot, M.** (2022). A panel of emerging EMT genes identified in malignant mesothelioma. *Sci Rep* **12**, 1007.

**Wu, Y., Liu, L., Shen, X., Liu, W. and Ma, R.** (2021). Plakophilin-2 Promotes Lung Adenocarcinoma Development via Enhancing Focal Adhesion and Epithelial-Mesenchymal Transition. *Cancer Manag Res* **13**, 559-570.

**Xiu, M. X., Liu, Y. M. and Kuang, B. H.** (2020). The oncogenic role of Jagged1/Notch signaling in cancer. *Biomed Pharmacother* **129**, 110416.

**Xu, N., Xiao, W., Meng, X., Li, W., Wang, X., Zhang, X. and Yang, H.** (2022a). Up-regulation of SLC27A2 suppresses the proliferation and invasion of renal cancer by down-regulating CDK3-mediated EMT. *Cell Death Discov* **8**, 351.

**Xu, Y., Gao, F., Zhang, J., Cai, P. and Xu, D.** (2022b). Fibroblast growth factor receptor 2 promotes the proliferation, migration, and invasion of ectopic stromal cells via activation of extracellular-signal-regulated kinase signaling pathway in endometriosis. *Bioengineered* **13**, 8360-8371.

**Yamamoto, H., Mukaisho, K., Sugihara, H., Hattori, T. and Asano, S.** (2011). Down-regulation of FXYD3 is induced by transforming growth factor-beta signaling via ZEB1/deltaEF1 in human mammary epithelial cells. *Biol Pharm Bull* **34**, 324-329.

**Yamashita, D., Minata, M., Ibrahim, A. N., Yamaguchi, S., Coviello, V., Bernstock, J. D., Harada, S., Cerione, R. A., Tannous, B. A., La Motta, C. and Nakano, I.** (2020). Identification of ALDH1A3 as a Viable Therapeutic Target in Breast Cancer Metastasis-Initiating Cells. *Mol Cancer Ther* **19**, 1134-1147.

**Ying, X., Sun, Y. and He, P.** (2015). Bone Morphogenetic Protein-7 Inhibits EMT-Associated Genes in Breast Cancer. *Cell Physiol Biochem* **37**, 1271-1278.

**Zeisberg, M., Hanai, J., Sugimoto, H., Mammoto, T., Charytan, D., Strutz, F. and Kalluri, R.** (2003). BMP-7 counteracts TGF-beta1-induced epithelial-to-mesenchymal transition and reverses chronic renal injury. *Nat Med* **9**, 964-968.

**Zeng, C., Long, J., Deng, C., Xie, L., Ma, H., Guo, Y., Liu, S. and Deng, M.** (2022). Genetic Alterations in Papillary Thyroid Carcinoma With Hashimoto(')s Thyroiditis： ANK3, an Indolent Maintainer of Papillary Thyroid Carcinoma. *Front Oncol* **12**, 894786.

**Zhai, Y., Iura, A., Yeasmin, S., Wiese, A. B., Wu, R., Feng, Y., Fearon, E. R. and Cho, K. R.** (2011). MSX2 is an oncogenic downstream target of activated WNT signaling in ovarian endometrioid adenocarcinoma. *Oncogene* **30**, 4152-4162.

**Zhang, M., Zhang, L., Geng, A., Li, X., Zhou, Y., Xu, L., Zeng, Y. A., Li, J. and Cai, C.** (2022). CDK14 inhibition reduces mammary stem cell activity and suppresses triple negative breast cancer progression. *Cell Rep* **40**, 111331.

**Zhang, N., Wang, F., Zhu, L., Chang, R., Mok, S. R. S., Peixoto, R. D., Tang, W. and Chen, Z.** (2023). Molecular mechanism of the miR-7/BCL2L1/P53 signaling axis regulating the progression of hepatocellular carcinoma. *Ann Transl Med* **11**, 12.

**Zhang, Y., Xia, M., Jin, K., Wang, S., Wei, H., Fan, C., Wu, Y., Li, X., Li, X., Li, G., et al.** (2018). Function of the c-Met receptor tyrosine kinase in carcinogenesis and associated therapeutic opportunities. *Mol Cancer* **17**, 45.

**Zheng, Y., Lu, J., Hu, X., Hu, X., Gao, X. and Zhou, J.** (2023). PRMT5/FGFR3/AKT Signaling Axis Facilitates Lung Cancer Cell Metastasis. *Technol Cancer Res Treat* **22**, 15330338231161139.

**Zhou, T., Luo, M., Cai, W., Zhou, S., Feng, D., Xu, C. and Wang, H.** (2018). Runt-Related Transcription Factor 1 (RUNX1) Promotes TGF-beta-Induced Renal Tubular Epithelial-to-Mesenchymal Transition (EMT) and Renal Fibrosis through the PI3K Subunit p110delta. *EBioMedicine* **31**, 217-225.

**Zhu, L. Y., Zhang, W. M., Yang, X. M., Cui, L., Li, J., Zhang, Y. L., Wang, Y. H., Ao, J. P., Ma, M. Z., Lu, H., et al.** (2015). Silencing of MICAL-L2 suppresses malignancy of ovarian cancer by inducing mesenchymal-epithelial transition. *Cancer Lett* **363**, 71-82.

**Zhu, X., Luo, X., Jiang, S. and Wang, H.** (2020). Bone Morphogenetic Protein 1 Targeting COL1A1 and COL1A2 to Regulate the Epithelial-Mesenchymal Transition Process of Colon Cancer SW620 Cells. *J Nanosci Nanotechnol* **20**, 1366-1374.

**Zuo, J., Ishikawa, T., Boutros, S., Xiao, Z., Humtsoe, J. O. and Kramer, R. H.** (2010). Bcl-2 overexpression induces a partial epithelial to mesenchymal transition and promotes squamous carcinoma cell invasion and metastasis. *Mol Cancer Res* **8**, 170-182.
